# D-Mannose treats obesity by increasing adipose Treg cells and suppressing gut *Firmicutes*

**DOI:** 10.64898/2026.08.10.743951

**Authors:** Peter Zanvit, Junji Xu, Nancy Guo, Dunfang Zhang, Michaela Prochazkova, Thierry Gauthier, Daxesh P. Patel, Alexander Cain, Wenwen Jin, Andrew Bynum, Frank J. Gonzalez, Yasmine Belkaid, WanJun Chen

## Abstract

Early-life microbiota represent an indispensable factor for the proper development and function of host metabolism and the immune system. We have demonstrated that neonatal exposure to antibiotics for the first 3 weeks (NeoATB) leads to obesity in adulthood, characterized by gut microbiota dysbiosis and dysregulated immune responses. Here, we demonstrate that feeding D-mannose suppresses NeoATB-induced obesity, accompanied by improved glucose tolerance and decreased insulin resistance. Mechanistically, D-mannose feeding decreased hypoxia and increased oxygenation and recovery of metabolic activity of adipocytes. D-mannose restored CD4^+^Foxp3^+^ST2^+^ Tregs, leading to a reduction of Th1 pro-inflammatory cells in the adipose tissue of NeoATB mice. Significantly, we revealed that D-mannose treatment reversed the dysregulated ratios of phylum *Firmicutes* to phylum *Bacteroidetes* in obese NeoATB mice, which was surprisingly attributed to D-mannose-mediated suppression of the growth of *Firmicutes* rather than an increase in the growth of *Bacteroidetes.* These findings should have therapeutic implications for the treatment of obesity in human patients.

## Introduction

Obesity represents a major health challenge as it substantially increases the risk of diseases such as type 2 diabetes, fatty liver disease, stroke, osteoarthritis, and several types of cancer (1). The dysbiosis of gut microbiota was identified as an important environmental factor that contributes to obesity (2), as transplantation of an “obese” microbiota into germ-free mice resulted in significantly increased adiposity compared with transplantation of “lean” microbiota (3, 4). Early-life perturbation of the gut microbiota is one of the most important factors contributing to obesity. In humans, early life microbiota perturbation using antibiotics (5, 6) or child delivery by Cesarean section (7, 8) has been associated with an increased risk of obesity development later in life. We have established a mouse model of obesity by treating mice with antibiotics only during the neonatal period (NeoATB, 0-21 days) and observed the development of obesity with metabolic abnormalities, including glucose intolerance and insulin resistance in the adult age (Zanvit et al., Barrier Immunity, in press) We have demonstrated that obesity in NeoATB mice is caused by a significantly reduced frequency and number of CD4^+^CD25^+^Foxp3^+^ regulatory T cells (Tregs), which leads to an increase in proinflammatory IFN-γ-producing T helper-1 (Th1) cells in visceral adipose tissue (VAT). In addition, elucidated that obesity is associated with and/or caused by a persistent dysbiosis of the gut microbiota, characterized by increased phylum *Firmicutes* and decreased phylum *Bacteroidetes*.

The success of D-mannose in inducing differentiation of Treg cells and suppressing experimental autoimmune type I diabetes and asthma in mice (9) has led us to hypothesize that oral feeding with D-mannose might ameliorate obesity in adult NeoATB mice. Thus, we treated the obese NeoATB mice with 20% D-mannose in drinking water. Indeed, D-mannose feeding had therapeutic effects on obesity by reducing the size of adipocytes and improving the metabolic activity of adipocytes in the VAT, which was attributed to increased Treg cells and decreased proinflammatory IFNγ-producing T cells in adipose tissues. Unexpectedly, D-mannose feeding corrected the dysregulated ratio of phylum *Firmicutes* to phylum *Bacteroidetes* in the gut of NeoATB obese mice. Additionally, D-mannose increased the abundance of phylum Verrucomicrobia, particularly the family *Verrucomicrobiaceae, in vivo*. Importantly, we revealed that D-mannose specifically suppressed the growth of the phylum *Firmicutes*, particularly the family *Ruminococcaceae*, rather than affecting the growth of *Bacteroidetes* members.

## Results

### D-mannose treatment ameliorates obesity in NeoATB mice

We have demonstrated that antibiotic treatment with vancomycin and polymyxin B during the first 3 weeks of life (neonatal age; NeoATB) leads to the development of obesity in adulthood (Zanvit et al., Barrier Immunity, in press). Recently, we showed that oral feeding of supraphysiological levels of D-mannose blocked the development of type I diabetes in NOD mice and reduced ovalbumin-induced airway inflammation in mice (9), which was mediated by an increase in CD4^+^CD25^+^Foxp3^+^ Treg cells. We thus hypothesized that D-mannose treatment might ameliorate obesity in adult NeoATB mice, as the obesity in these mice was attributed to the reduction of Treg cells in the fat tissue. To test this hypothesis, we treated one-year-old normal (control, CTRL) and obese NeoATB mice with 20% D-mannose in drinking water or water alone for 4 weeks (**Supplemental Figure 1A, B**). Surprisingly, we observed a significant reduction in body weight in the D-mannose-treated obese NeoATB mice, whereas no significant effect was observed in the D-mannose-treated control mice (**Figure 1A**). There was no significant difference in the food intake among all groups (**Supplemental Figure 1C**). Consistently, both the size and weight of visceral adipose tissues (VAT) in D-mannose-treated NeoATB mice were significantly reduced. In contrast, no significant difference was observed in the size and weight of fat in control mice treated with D-mannose (**Figure 1B, C**). Histological analysis of VAT revealed a significant decrease in adipocyte size in D-mannose-treated NeoATB obese mice, but not in control mice (**Figure 1D, E**). Obesity is associated with a spectrum of liver abnormalities, known as non-alcoholic fatty liver disease (NAFLD), characterized by an increase in intrahepatic triglyceride content (steatosis) (10). Indeed, in the NeoATB mice, but not in control mice, we observed significant ballooning in the liver, a clear induction of steatosis and NAFLD development (**Figure 1F, G**). However, after treatment with D-mannose, NeoATB obese mice showed substantial improvement in hepatic steatosis (**Figure 1F, G**). Next, we investigated the effect of D-mannose treatment on metabolism in obese NeoATB mice. Using the glucose tolerance test, we found apparent glucose intolerance in obese NeoATB mice, which was significantly improved by D-mannose treatment (**Figure 1H, Supplemental Figure 1D**). Surprisingly, D-mannose treatment also elevated glucose response in control mice (**Figure 1H, Supplemental Figure 1D**). Consistently, D-mannose treatment also significantly improved insulin resistance in both control and NeoATB mice (**Figure 1I, Supplemental Figure 1E**).

**Figure 1.**
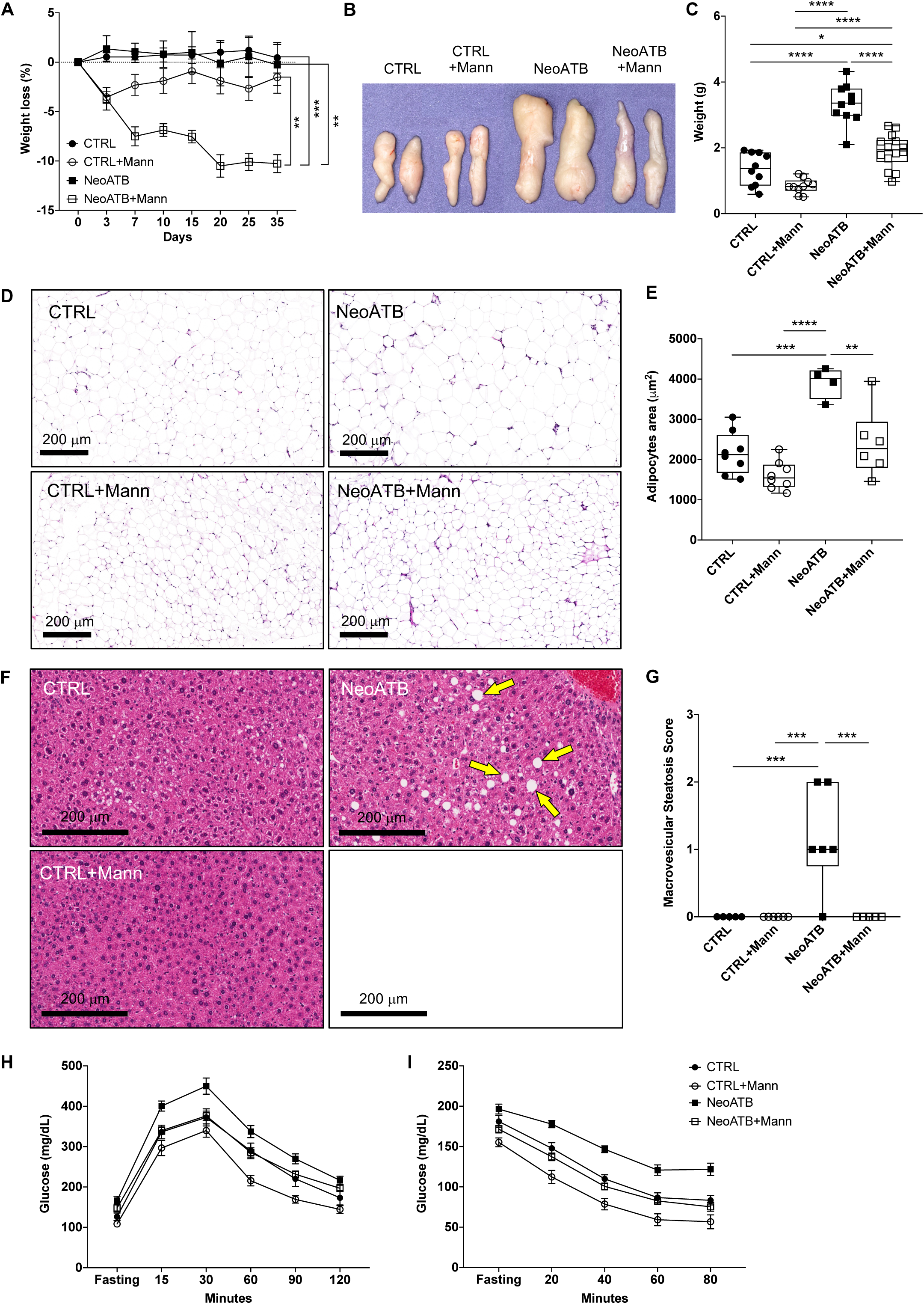
D-mannose suppresses obesity in obese NeoATB mice. A. Aged Control or obese NeoATB mice were treated with 20% D-mannose (CTRL+Mann and NeoATB+Mann) or left untreated (CTRL and NeoATB). Representative figure showing weight loss among different groups. B. Representative photograph of visceral adipose tissue among different groups. C. Summarizing data showing the weight of visceral adipose tissue among different groups. D. Histology (H&E) staining of visceral adipose tissue among different groups. The scale bar represents 200μm. E. Adipocyte area was determined using AdipoCount software among different groups. F. Histology (H&E) staining of liver tissue among different groups. Yellow arrows indicate fat accumulation in the liver. G. Macrovesicular steatosis score among different groups. The macrovesicular score was performed blindly by two independent investigators. H. Glucose tolerance test. Mice were fasted overnight (15-16 h) and intraperitoneally injected with 2 g/kg glucose. Blood glucose levels were measured in intervals of 0, 15, 30, 60, 90, and 120 min. I. Insulin tolerance test. Mice were fasted for 4-6 h and injected intraperitoneally with 0.75 U/kg of insulin. Blood glucose levels were measured in intervals of 0, 20, 40, 60, and 80 min. Data are representative of 2-3 independent experiments. Statistical analysis was determined using one-way ANOVA (* p<0.05, ** p<0.01, *** p<0.001, **** p<0.0001). Each point in the figure represents an individual animal.

To further validate the positive effects of D-mannose treatment on obesity and metabolism, we employed the high-fat diet (HFD)-induced obesity model. We fed mice a HFD continuously from 3 through 30 weeks of age and treated the HFD-fed mice with D-mannose (20% in water, HFD+Mann) or water alone (HFD) from 18 weeks through 30 weeks of age (**Supplemental Figure 2A**). Similar to NeoATB mice, D-mannose treatment significantly lowered body weight and reduced the weight and size of visceral fat in HFD-fed mice (**Supplemental Figure 2C**). D-Mannose-treated HFD mice exhibited reduced size of adipocytes (**Supplemental Figure 2D, E**). Moreover, we tested the effect of D-mannose on obesity development in young NeoATB mice. In this study, young NeoATB mice were treated with D-Mannose or left untreated. Body weight was monitored for over 200 days. Our data showed that D-mannose treatment could reduce obesity in adult NeoATB mice **(Supplemental Figure 3A)**. Additionally, we conducted a fecal microbiota transfer (FMT) experiment. In this experiment, fecal pellets were collected from either adult NeoATB or NeoATB D-Mannose-treated mice and given to 20-week-old NeoATB mice via gavage. Our results indicate that transferring microbiota from D-Mannose-treated mice can indeed help prevent obesity in NeoATB mice **(Supplemental Figure 3B)**.

Collectively, these data revealed that D-mannose treatment suppresses obesity and improves metabolism.

### D-Mannose corrects dysregulated metabolism in adipocytes of obese mice

We next investigated the effects of D-mannose treatment on metabolism in adipocytes, with a focus on the insulin response in NeoATB mice. Insulin exerts all of its known physiological effects by binding to the insulin receptor (INSR) on the plasma membrane of target cells (11). We first examined the expression of mRNA and protein levels of the insulin receptor in VAT from control and NeoATB mice after treatment with D-mannose. The insulin receptor (*Insr*) mRNA was significantly reduced in the VAT of NeoATB mice compared to control mice, as expected, which was, however, significantly improved after D-mannose treatment (**Figure 2A**). Interestingly, D-mannose feeding failed to significantly change *Insr* mRNA levels in control mice (**Figure 2A**). Consistent with the mRNA expression, western blot analysis revealed a reduced INSR protein level in the VAT of NeoATB mice, which was restored by D-mannose treatment (**Figure 2B**). Besides the insulin receptor, insulin increases glucose uptake into fat cells through the regulated trafficking of vesicles that contain glucose transporter type 4 (SLC2A4) (12). The defects in the glucose uptake represent an early step in the development of type 2 diabetes. Thus, we examined the expression of *Slc2a4* mRNA in the fat of obese NeoATB mice and found that NeoATB VAT exhibited significantly decreased expression and SLC2A4 protein levels compared to control mice (**Figure 2C, D**); D-mannose treatment however, reversed this decrease (**Figure 2C, D**). Thus, D-mannose corrected the insulin resistance found in NeoATB mice.

**Figure 2.**
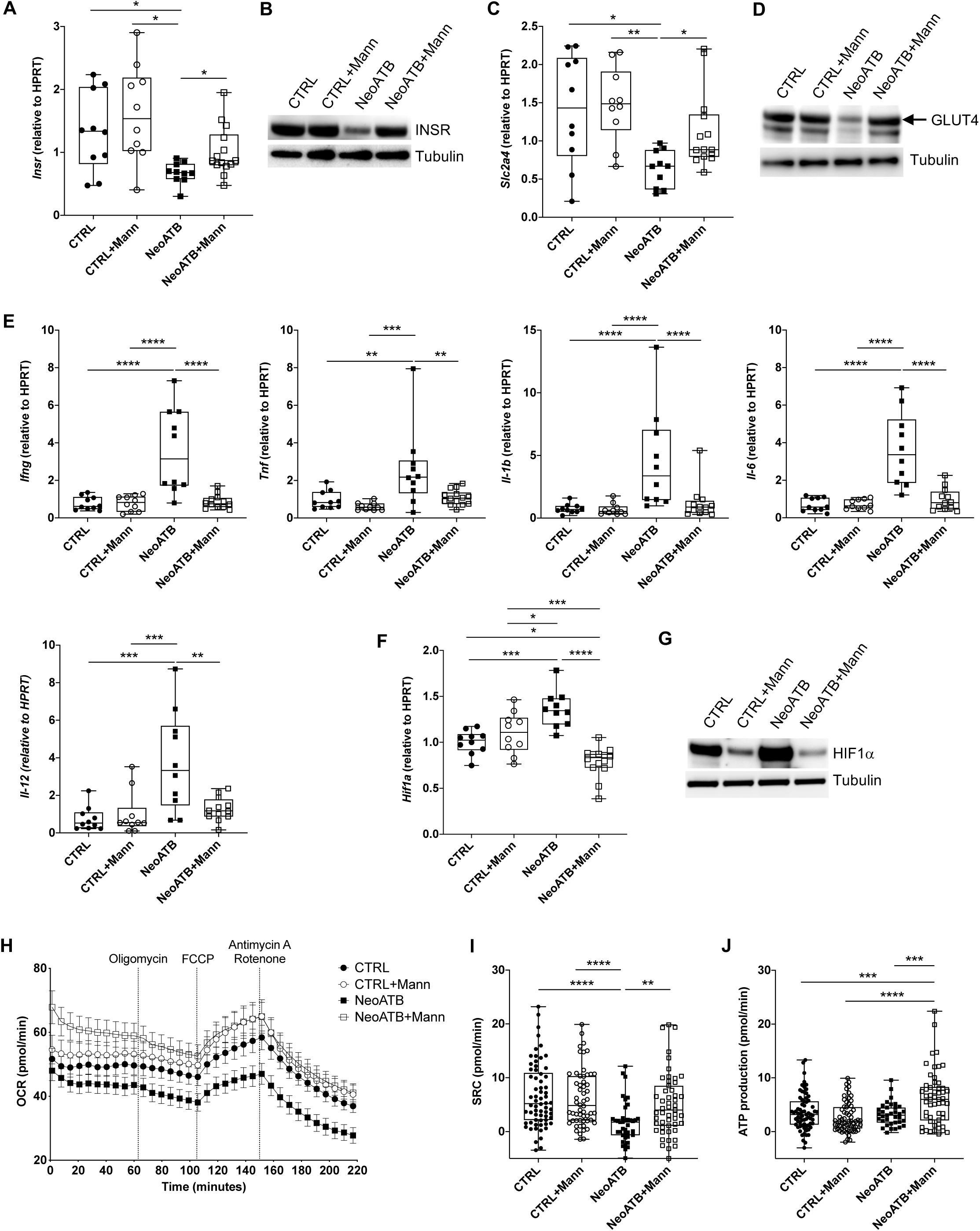
D-mannose improves the metabolism of adipose tissue in NeoATB mice. A. Expression of insulin receptor (*Insr*) mRNA in adipose tissue of Control, NeoATB or D-mannose-treated Control or NeoATB mice. B. Western blot of insulin receptor protein in the adipose tissue of Control, NeoATB, or D-mannose-treated Control or NeoATB mice. C. Expression of *Slc2a4* (encoding GLUT4) mRNA in adipose tissue of Control, NeoATB, or D-mannose-treated Control or NeoATB mice. D. Western blot of GLUT4 protein in adipose tissue of Control, NeoATB or D-mannose-treated Control or NeoATB mice. E. Expression of pro-inflammatory cytokine mRNAs in adipose tissue of Control, NeoATB, or D-mannose-treated Control or NeoATB mice. F. Expression of hypoxia inducible factor 1 (*Hif1a*) mRNA in the adipose tissue of Control, NeoATB, or D-mannose treated Control or NeoATB mice. G. Western blot of HIF1A protein in adipose tissue of Control, NeoATB or D-mannose-treated Control or NeoATB mice. H. Spheroid 3D tissue culture, showing oxygen consumption rate in the adipose tissue of Control, NeoATB, or D-mannose-treated Control or NeoATB mice. Data are representative of 8 CTRL, 8 CTRL+Mann, 6 NeoATB, and 7 NeoATB+Mann mice. I. Spare respiratory capacity among different groups. Data are representative of 8 CTRL, 8 CTRL+Mann, 6 NeoATB, and 7 NeoATB+Mann mice. Data are shown as SRC per treatment. 8 pieces of visceral adipose tissue (weight range 5-8 mg) were tested per mouse. J. ATP production among different groups measured using the Seahorse instrument. All data are representative of 2 independent experiments. Statistical analysis was determined using one-way ANOVA (* p<0.05, ** p<0.01, *** p<0.001, **** p<0.0001). Each point in the figure represents an individual animal.

Obesity is associated with low-grade inflammation in adipose tissue and increased expression of inflammatory cytokines such as IFNγ, TNF, IL-6, and IL1β, which were shown to potentiate insulin resistance in part via suppression of the insulin receptor (13–16). We found that the mRNA levels encoding IFNγ, TNF, IL1, β, IL-6, and IL-12 were all significantly increased in the VAT of obese NeoATB mice, and D-mannose treatment eliminated the increase in these pro-inflammatory cytokines in NeoATB mice (**Figure 2E**). Recent reports suggest that hypoxia is a risk factor for chronic inflammation in adipose tissue (17, 18). We thus determined that the mRNA expression and protein levels of hypoxia-inducible factor 1 alpha (HIF1α) were significantly increased in VAT of NeoATB mice, which were completely reversed by D-mannose treatment (**Figure 2F, G**). Interestingly, the levels of HIF1α in D-mannose-treated NeoATB mice were even lower than those in control mice (**Figure 2F**). Of note, D-mannose also reduced the HIF1A protein in the VAT of control mice (**Figure 2G**). To further validate the hypoxia, we directly tested oxygen levels in adipose tissue by using the Seahorse spheroid 3D culture system (**Figure 2H**). Adipose tissue respiration was determined by observing how the oxygen consumption rate (OCR) changes in response to drugs that modulate mitochondrial activity. We detected reduced respiration of VAT in NeoATB mice compared to control mice, while D-mannose treatment completely reversed this reduction (**Figure 2H**). Consistent with increased HIF1α, obese NeoATB mice exhibited a substantial decrease in spare respiratory capacity (SRC) of adipose tissue, which was again completely restored by D-mannose treatment (**Figure 2I**). Finally, D-mannose treatment of NeoATB mice also significantly increased ATP production in adipose tissue (**Figure 2J**). Taken altogether, D-mannose treatment corrects hypoxia by improving oxygenation of adipocytes, which helps to suppress inflammation in fat tissue and consequently reduces insulin resistance of adipocytes.

### D-Mannose suppresses obesity by increasing adipose ST2^+^ regulatory T cells

As fat CD4^+^Foxp3^+^ regulatory T cells (Treg) are crucial in controlling inflammation and metabolism in adipose tissue (19) and obese NeoATB mice exhibited a significant decrease in adipose Tregs, we hypothesized that D-mannose could restore Tregs in the adipose tissue of NeoATB mice. Indeed, D-mannose feeding significantly increased the frequency and total number of CD4^+^CD25^+^Foxp3^+^ Treg cells in the VAT in NeoATB mice, which was almost completely recovered to the levels of control mice (**Figure 3A, B**). Interestingly, D-mannose feeding also significantly increased the frequency, but not the absolute number, of VAT Treg cells in control mice (**Figure 3A, B**). It was known that fat Tregs expressing ST2 (IL-33 receptor) are crucial in the regulation of inflammation in adipose tissue (20). We found significantly decreased frequency and total number of CD4^+^CD25^+^Foxp3^+^ST2^+^ Tregs (ST2^+^ Tregs) in the VAT of NeoATB mice compared to control mice (**Figure 3A, C**). D-mannose treatment reversed this defect in ST2^+^Tregs in the VAT of NeoATB mice (**Figure 3A, C**). Of note, no significant difference in ST2^+^ Tregs between untreated and D-mannose-treated control mice was found (**Figure 3A-C**).

**Figure 3.**
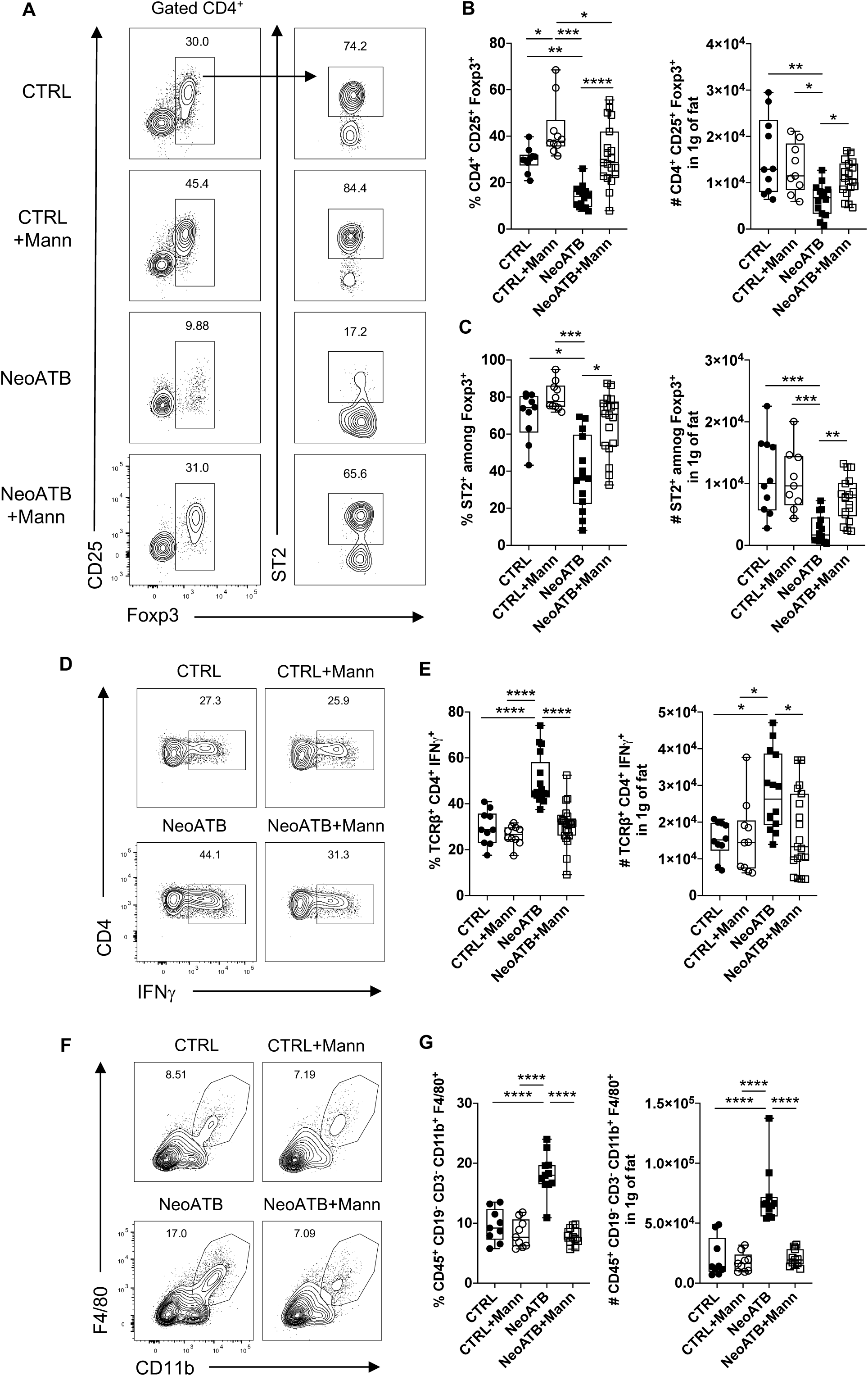
D-mannose treatment increases adipose regulatory T cells. A. Representative FACS plots showing frequency of adipose regulatory T (Treg) cells among different groups. Cells were gated as CD45^+^TCRβ^+^CD4^+^CD25^+^Foxp3^+^ or CD45^+^TCRβ^+^CD4^+^CD25^+^Foxp3^+^ST2^+^. B. Summarizing figure showing frequency and total number of CD45^+^TCRβ^+^CD4^+^CD25^+^Foxp3^+^ cells among different groups. C. Summarizing figure showing frequency and total number of CD45^+^TCRβ^+^CD4^+^CD25^+^Foxp3^+^ST2^+^ cells among different groups. D. Representative FACS plots showing frequency of adipose Th1 CD4^+^IFNγ^+^ cells among different groups. E. Summarizing figure showing frequency and total number of adipose Th1 CD4^+^IFNγ^+^ cells among different groups. F. Representative FACS plots showing frequency of adipose macrophages (Gated as: CD45^+^CD19^-^CD3^-^CD11b^+^F4/80^+^) among different groups. G. Summarizing data showing frequency and total number of adipose infiltrating macrophages among different groups. Data are pooled from 3 independent experiments. Statistical analysis was determined using one-way ANOVA (* p<0.05, ** p<0.01, *** p<0.001, **** p<0.0001). Each point in the figure represents an individual animal.

We next investigated how D-mannose upregulated adipose Treg cells. We first observed reduced proliferation of CD4^+^Foxp3^+^ Treg cells in the VAT of obese NeoATB mice, as indicated by decreased Ki67 expression compared to control mice **(Supplemental Figure 4A)**. D-Mannose treatment, however, significantly increased Ki67^+^ Treg cells in NeoATB mice (**Supplemental Figure 4A**), suggesting D-mannose promotion of ST2^+^ Treg expansion in the VAT.

Consistent with the increased proliferation of CD4^+^Foxp3^+^ST2^+^ Treg cells, D-Mannose treatment increased VAT expression *of il-33* **(Supplemental Figure 4C).** Moreover, we have confirmed increased frequency of VAT CD45^-^CD31^-^SCA1^+^PDGFR^+^ VAT mesenchymal stromal cells **(Supplemental Figure 4D)**, which were identified as a major source of IL-33 (Spallanzani et al. 2019).

In addition to the increase in Treg cell proliferation, D-mannose treatment also significantly increased expression of mRNAs encoding TGFβ receptor I and II in adipose naïve CD4^+^CD25^-^ T cells of NeoATB mice **(Supplemental Figure 4E, F)**, suggesting that an increased capacity of local conversion of Treg cells from naïve T cells, as TGFβ receptor I-mediated signaling is essential for Treg conversion (21). Although D-mannose treatment did not change the total levels of VAT TGFβ protein **(Supplemental Figure 4G)**, it may enhance the activation of latent TGFβ in VAT (Zhang et al, 2017).

As a crucial function of adipose Tregs (especially ST2^+^ Tregs) is to control inflammation of adipose tissue (19), the reduction of Treg numbers in adipose tissue indeed led to a significant increase in the proliferation of CD4^+^Foxp3^-^ responder T cells in obese NeoATB mice (**Supplemental Figure 4B**). D-Mannose treatment, however, significantly decreased the CD4^+^Foxp3^-^Ki67^+^ responder T cells in NeoATB mice (**Supplemental Figure 4B**). In addition to the decrease in T responder cell proliferation, D-mannose treatment also significantly inhibited the accumulation of pro-inflammatory immune cells such as CD4^+^IFN-γ^+^ Th1 cells (**Figure 3D, E**), TCRγ8^+^IFNγ^+^ **(Supplemental Figure 5A)** T cells, and M1-like proinflammatory macrophages in the VAT of NeoATB mice (**Figure 3F, G**). In contrast, D-mannose treatment slightly increased the number of CD4^+^IL-5^+^ Th2 cells in the VAT of NeoATB mice **(Supplemental Figure 5E)**. Interestingly, there were no significant differences in the frequency or total number of CD4^+^IL-10^+^, CD8^+^IFNγ^+^ cells, and Th17 or TCRγ8^+^IL-17^+^ T cells in the adipose tissue among all groups **(Supplemental Figure 5)**. To determine if the D-mannose-mediated increase in adipose Treg cells was indeed responsible for the amelioration of obesity in NeoATB mice, we depleted Treg cells using anti-CD25 antibody treatment in D-mannose-treated mice **(Supplemental Figure 6).** As in the adipose tissue, where more than 70% of adipose Tregs are ST2^+^ CD25^+^ **(Figure 3, Supplemental Figure 6A**), we found that anti-CD25 antibody treatment indeed significantly depleted ST2^+^Foxp3^+^ Tregs in the VAT (**Supplemental Figure 6D-E**). The depletion of adipose ST2^+^ Treg cells abolished the suppressive function of D-mannose treatment in obesity and metabolic syndrome **(Supplemental Figure 6D-E)**, which was attributed to the failure in decreasing Th1 cells in the VAT of NeoATB mice **(Supplemental Figure 6F-G)**. As expected, anti-CD25 antibody injection did not affect the numbers of VAT macrophages in D-mannose-treated NeoATB mice **(Supplemental Figure 5H, I)**. These data indicate that the reduction of obesity was attributed to upregulation of ST2^+^ Treg cells in adipose tissue.

### D-Mannose feeding increases adipose group 2 innate lymphoid cells

In addition to Treg cells, group 2 innate lymphoid cells (ILC2s) in murine adipose tissue also act to control obesity (22). Adipose ILC2s express Gata3, Stem Cells Antigen-1 (SCA-1) and ST2 and produce cytokines such as IL-5 and IL-13 (23). In obese NeoATB mice, we found a significantly reduced frequency and absolute number of VAT ILC2s (CD45^+^Lin^-^Gata3^+^ST2^+^SCA-1^+^) (**Supplemental Figure 7A, B, C**). D-Mannose feeding significantly upregulated the frequency and total number of ILC2 cells in the VAT of NeoATB mice (**Supplemental Figure 7A, B, C**). Moreover, the frequency and total number of IL-5^+^ and IL-13^+^ ILC2s were significantly decreased in obese NeoATB mice, and this decrease was restored by D-mannose treatment (**Supplemental Figure 7A, D, E**). However, D-mannose treatment did not significantly influence the numbers of ILC2s or their cytokine production in the VAT of control mice (**Supplemental Figure 7**). Thus, D-mannose also increases ILC2 cells in the adipose tissue of the obese NeoATB mice, suggesting a role for ILC2 in D-mannose-mediated suppression of obesity.

### D-Mannose feeding corrects dysbiosis of gut microbiota in obese mice

In addition to Tregs in fat tissue, the gut microbiota plays a crucial role in regulating the immune response and metabolism. Dysbiosis of the microbiota has been shown in both animal models of obesity and obese humans (4, 24–26). As D-mannose treatment was performed orally in drinking water, we reasoned whether the anti-obesity effect of D-mannose could be mediated by modifying the gut microbiome. For this, we first measured the total bacterial load in the feces of control and NeoATB mice with or without D-mannose treatment. Compared to the control groups, D-mannose-treated NeoATB mice had significantly increased bacterial load in their fecal pellets, as indicated by the higher number of 16S rDNA copies (**Figure 4A**). However, 16S sequencing analysis did not reveal significant changes in the alpha-diversity (Observed species Index and Simpson Index) among all groups (**Figure 4B**). Beta diversity analysis (Bray-Curtis Index) revealed discrete clustering between control, D-mannose-treated control, NeoATB, and D-mannose-treated NeoATB mice (**Figure 4C**). On the phylum level, as expected, phylum *Firmicutes* exhibited significantly greater abundance in the feces of obese NeoATB mice, whereas the abundance of phylum *Bacteroidetes* in the same mice was significantly reduced when compared to that of control mice (**Figure 4D**). However, D-mannose treatment significantly reduced the abundance of phylum *Firmicutes* and slightly increased the abundance of phylum *Bacteroidetes* in obese NeoATB mice. Interestingly, D-mannose treatment also significantly increased the abundance of phylum *Verrucomicrobia* in both control and NeoATB mice (**Figure 4D, E**). At the family level, D-mannose treatment significantly decreased the abundance of families: *Bacteroidaceae* and *Rikenellaceae* (phylum *Bacteroidetes*), as well as *Ruminococcaceae*, *Christensenellaceae*, *Clostridiaceae,* and *Mogibacteriaceae* (phylum Firmicutes), but increased the abundance of families *S24-7* (phylum *Bacteroidetes*) and *Verrucomicrobiaceae* (phylum *Verrucomicrobia*) in obese NeoATB mice (**Figure 4F, G**). Intriguingly, D-mannose treatment of control mice significantly reduced the abundance of family *Mogibacteriaceae* (phylum Firmicutes) and increased the abundance of family *Verrucomicrobiaceae* (phylum *Verrucomicrobia*) compared to untreated control mice (**Figure 4F, G**). To investigate the relationship between CTRL, NeoATB or D-mannose-treated CTRL and NeoATB gut microbiome functions, we predicted the potential metagenomes from the community profiles of randomized 16S rRNA genes using PICRUSt (27). The inferred gene families were annotated and combined with level 3 pathways (KEGG) for statistical testing and evaluation using STAMP software (28). Using PICRUSt (KEGG, level 3), we identified over 100 significant metabolic pathways where the difference in % of relative frequency was significant among the four groups **(Supplemental Figure 8A)**. D-Mannose treatment of NeoATB mice significantly corrected the following metabolic pathways (KEGG, level 3): ABC transporters, phosphotransferase system (PTS), chaperones and folding catalysts, transcription factors, citrate cycle, pentose phosphate pathway, oxidative phosphorylation, tyrosine and tryptophan biosynthesis, biotin metabolism and folate biosynthesis. On the other hand, D-mannose treatment significantly reduced bacterial chemotaxis and flagellar assembly and significantly increased bacterial secretion system, arginine and proline metabolism, tryptophan metabolism, and glycosyltransferases metabolism in both CTRL and NeoATB-treated mice **(Supplemental Figure 8B)**.

**Figure 4.**
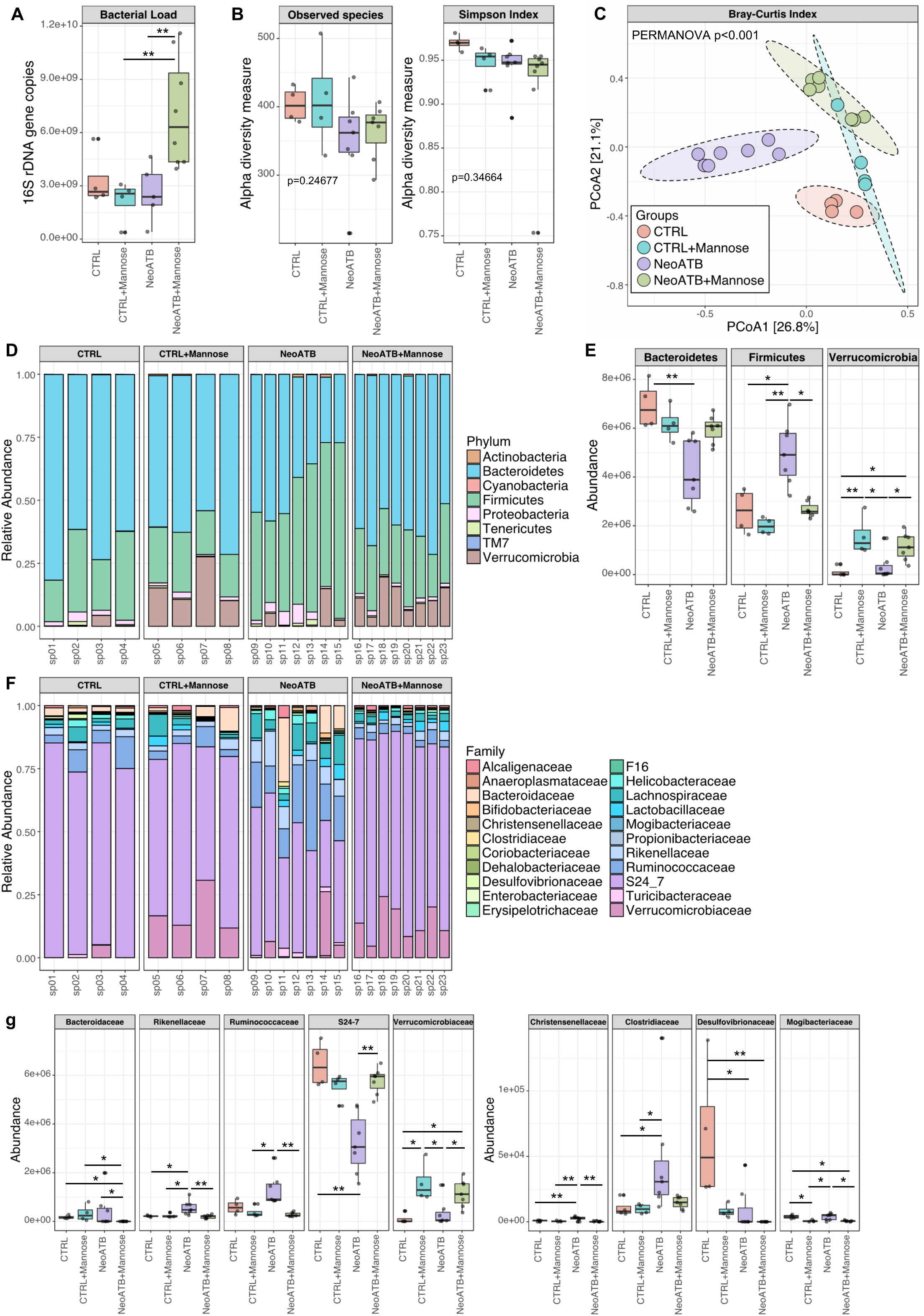
D-mannose treatment modifies the gut microbiome of NeoATB mice. A. Total bacterial load determined as the number of 16S rDNA gene copies using qPCR among different groups. B. Alpha diversity measure (Observed species, Simpson Index) among different groups. C. Beta diversity measure (Bray-Curtis Index) among different groups. D. Phylum-level taxa abundance among different groups. E. Summarizing plot showing significant changes of the microbiota on the phylum level among different groups. F. Family-level taxa abundance among different groups. G. Summarizing plots showing significant changes in the microbiota on the family level among different groups. Data are pooled from 2 independent experiments. Statistical analysis was determined using the Kruskal-Wallis test (* p<0.05, ** p<0.01). Each point in the figure represents an individual sample.

We then extended our analysis of the D-mannose effects on the gut microbiota in the HFD-induced obesity model that is also characterized by a decrease of *Bacteroidetes* and an increase of *Firmicutes*, two alterations that have been associated with obesity (26). First, we determined that D-mannose-treated HFD-fed mice displayed significantly increased bacterial load in feces compared to untreated HFD-fed mice (**Supplemental Figure 9A**). 16S sequencing analysis revealed reduced alpha-diversity (Observed species Index and Shannon index) in D-mannose-treated HFD-fed mice (p=0.036209 and p=0.019575, respectively) (**Supplemental Figure 9B**). Beta diversity (Bray-Curtis Index) revealed discrete clustering in the gut microbiome between untreated HFD-fed and D-mannose-treated HFD-fed mice (**Supplemental Figure 9C**). At the phylum level, Linear discriminant analysis (LDA=3; p<0.05) revealed a significant decline in the abundance of *Firmicutes* and *Proteobacteria* and a significant increase in the abundance of *Verrucomicrobia* in the feces of D-mannose-treated HFD-fed mice (**Supplemental Figure 9D, F**). Consistent with the D-mannose effects in NeoATB mice (**Fig. 4**), at the family level, D-mannose treatment significantly reduced family *Ruminococcaceae*, *Lactobacillaceae*, *Clostridiaceae*, *Streptococcaceae*, *Peptococcaceae,* and *Mogibacteriaceae* (phylum *Firmicutes*) and *Desulfovibrionaceae* (phylum *Proteobacteria*), and significantly increased the abundance of family Verrucomicrobiaceae (phylum *Verrucomicrobia*) in the feces of HFD-fed mice (**Supplemental Figure 9E, G**). These results are quite similar to the microbiota profile in D-mannose-treated NeoATB mice (**Figure 4**). At the species level, we found that the abundance of *Ruminococcus gnavus* was significantly reduced, but *Akkermansia muciniphila* was enriched by D-mannose treatment in HFD-fed mice (**Supplemental Figure 9H**). Collectively, these data revealed a significant effect of D-mannose treatment on the gut microbiota, specifically by suppressing the phylum Firmicutes and increasing the phylum *Verrucomicrobia* in both NeoATB and HFD-fed models of obesity.

### D-Mannose feeding reduces propionate in the gut of obese NeoATB mice

Despite intake of the same diet (in terms of quantity and composition of fiber), differences in gut microbiota composition can lead to different non-digestible carbohydrate fermentation profiles and varied production of short-chain fatty acids (SCFAs). Among the three types of SCFAs, i.e., acetate, butyrate, and propionate, the increase in propionate was positively associated with obesity. For instance, when compared with lean mice, mice with genetically-induced obesity have an increased ratio of *Firmicutes* to *Bacteroidetes*, which has been associated with increased cecal concentrations of acetate and butyrate in lean or acetate and propionate in obese mice, respectively (4, 29). In humans, an increase in the fecal concentrations of propionate and a decreased ratio of *Bacteroidetes* to *Firmicutes* were reported in obese individuals compared with lean individuals (30, 31). As obese NeoATB mice showed increased ratios of *Firmicutes* to *Bacteroidetes* (**Figure 4**), which was reversed by D-mannose treatment (**Figure 4**), we hypothesized that D-mannose might also change the concentrations of SCFA, especially a decrease in propionate in obese NeoATB mice. For this, we measured the levels of fecal SCFA and found that obese NeoATB mice indeed had significantly increased concentrations of propionate, acetate, and butyrate compared to control mice (**Figure 5A, B, C**). Of note, however, among the three SCFAs, propionate levels remained the highest at ∼50-100 mM/mg, compared to ∼2 mM/mg of acetate and nM/mg of butyrate (**Figure 5A, B, C**). Strikingly, D-mannose treatment significantly reduced the levels of propionate in the feces of NeoATB mice (**Figure 5C**). D-Mannose also decreased the amounts of butyrate, but not acetate (**Figure 5A-B**). The effect of propionate in promoting obesity requires expression of fatty acid binding protein 4 (FABP4), as mice deficient for *Fabp4* were protected from the development of obesity induced by propionate (32). We next measured *Fabp4* mRNA and found that it was significantly increased in the VAT of NeoATB mice (**Figure 5D**). Intriguingly, D-mannose treatment significantly decreased the levels of *Fabp4* mRNA in NeoATB mice (**Figure 5D**). To test the effect of propionate on reducing obesity in NeoATB mice, we treated 3-week-old NeoATB mice with D-mannose or D-mannose plus 1% Propionate. Mice’s body weights were monitored for over 200 days. At day 216, the mice were harvested, and the weight of visceral adipose tissue (VAT) was measured. Our data indicate that Propionate effectively counteracted the effects mediated by D-Mannose. Mice treated with the D-Mannose plus Propionate combination displayed significantly higher body weight and greater adipose tissue weight **(Figure 5E)**. SCFAs may also affect host metabolism by activating G-protein-coupled cell surface receptors, specifically GPR41 (also known as free fatty acid receptor 3) and GPR43 (also known as free fatty acid receptor 2) (33). In contrast to FABP4, however, we did not find a significant difference in expression of *Ffar2* (GPR43) or *Ffar3* (GPR41) in the VAT of NeoATB mice compared to control mice (**Figure 5F-G**), and D-mannose treatment did not significantly change the levels of *Ffar2* and *Ffar3*. These data highlight a role for D-mannose in decreasing Propionate in the gut of NeoATB mice, further validating the function of D-mannose in restoring the balance of gut microbiota.

**Figure 5.**
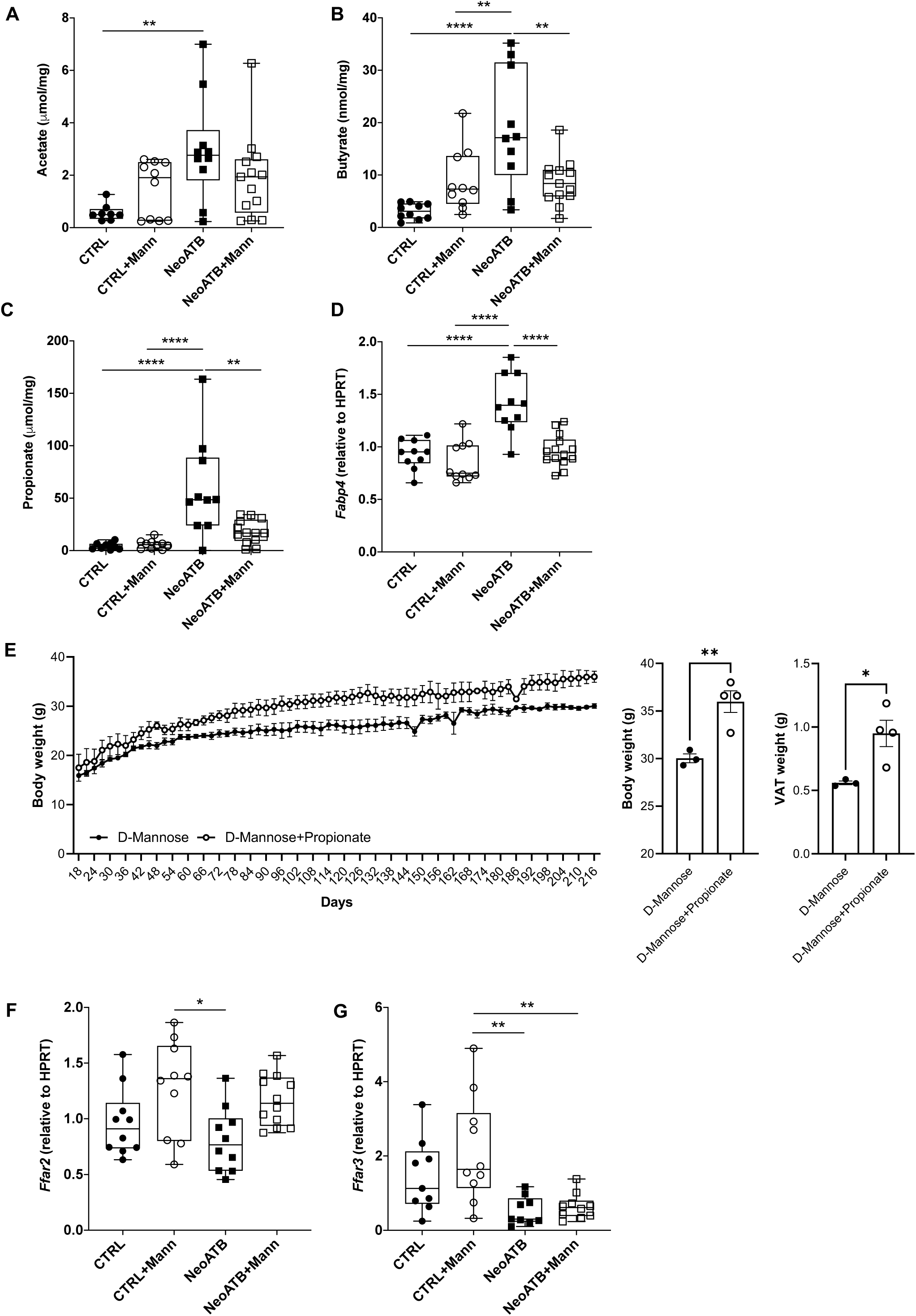
D-mannose treatment reduces propionate levels in NeoATB mice. A. Summarizing plots showing levels of the short-chain fatty acid acetate in the feces of NeoATB mice determined by GS-MS analysis. Data is shown as μmol/mg of fecal content. B. Summarizing plots showing levels of the short-chain fatty acid butyrate in the feces of NeoATB mice determined by GS-MS analysis. Data is shown as nmol/mg of fecal content. C. Summarizing plots showing levels of the short-chain fatty acid propionate in the feces of NeoATB mice determined by GS-MS analysis. Data is shown as μmol/mg of fecal content. D. Summarizing plot showing levels of adipose *Fabp4* mRNA among different groups. E. Summarizing data showing body weight time course of 3-week-old NeoATB mice, treated with D-Mannose or a combination of D-Mannose and 1% Propionate. Bar graphs showing Body weight and the weight of VAT measured at the end of the experiment. F. Summarizing plot showing levels of adipose *Ffar2* mRNA (encoding GPR43) among different groups. G. Summarizing plot showing levels of adipose *Ffar3* mRNA (encoding GPR41) among different groups. Data are pooled from 2 independent experiments. Statistical analysis was determined using one-way ANOVA (* p<0.05, ** p<0.01, *** p<0.001, **** p<0.0001). Each point in the figure represents an individual animal. Data in E are representative of 1 experiment, and statistical analysis was performed using a two-tailed *t-test* (* p<0.05, ** p<0.01).

#### D-Mannose directly suppresses the growth of *Firmicutes*

Our findings that D-mannose treatment reduced the abundance of *Firmicutes* and increased the abundance of *Verrucomicrobia* and *Bacteroidetes* in the gut of obese NeoATB mice (**Figure 4, Suplemental Figure 10**) led us to hypothesize that D-mannose might regulate the growth of specific microbiota over others. For this, we selected culturable candidates from *Firmicutes (Ruminococcus gnavus, Lachnospiraceae bacterium), Bacteroidetes (Bacteroides acidifaciens, Bacteroides ovatus),* and *Verrucomicrobia (Akkermansia muciniphila)* to test the effect of D-mannose on the growth of particular bacteria directly in culture. We first selected minimal media that could support the growth of the selected bacteria under anaerobic conditions. All of the bacteria were able to grow in heart infusion media without the addition of dextrose. As glucose is known to support the growth of many bacteria, we selected glucose as a control sugar. Thus, the selected bacteria were cultured either without sugar (no carbon source), or with 0.5% D-glucose or 0.5% D-mannose. Unexpectedly, while 0.5% D-glucose promoted vigorous growth of both *Ruminococcus gnavus* and *Lachnospiraceae bacterium* (*Firmicutes*) compared to those in no carbon source media, 0.5% D-mannose failed to enhance the growth of these two bacteria in culture (**Figure 6A, B**). Addition of 0.5% D-glucose into 0.5% D-mannose media partially restored the growth of these two bacteria suppressed by D-mannose in the culture (**Figure 6A, B**). Further experiments revealed that higher ratios of D-mannose:D-glucose more efficiently inhibited growth of *Ruminococcus gnavus* and *Lachnospiraceae bacterium* (**Supplemental Figure 10A-B**). In marked contrast, *Bacteroides acidifaciens* and *Bacteroides ovatus* (*Bacteroidetes*) as well as *Akkermansia muciniphila* (*Verrucomicrobia*) grew similar in either 0.5% D-glucose- or 0.5% D-mannose-supplemented media than under no carbon source media (**Figure 6C, D, E**). The data indicate that D-mannose specifically suppresses the growth of certain bacteria of phylum *Firmicutes*, which reveal a mechanism for the regulatory function of D-mannose treatment in rebalancing the ratio between phylum *Firmicutes* and *Bacteroidetes or Verrucomicrobia*.

**Figure 6.**
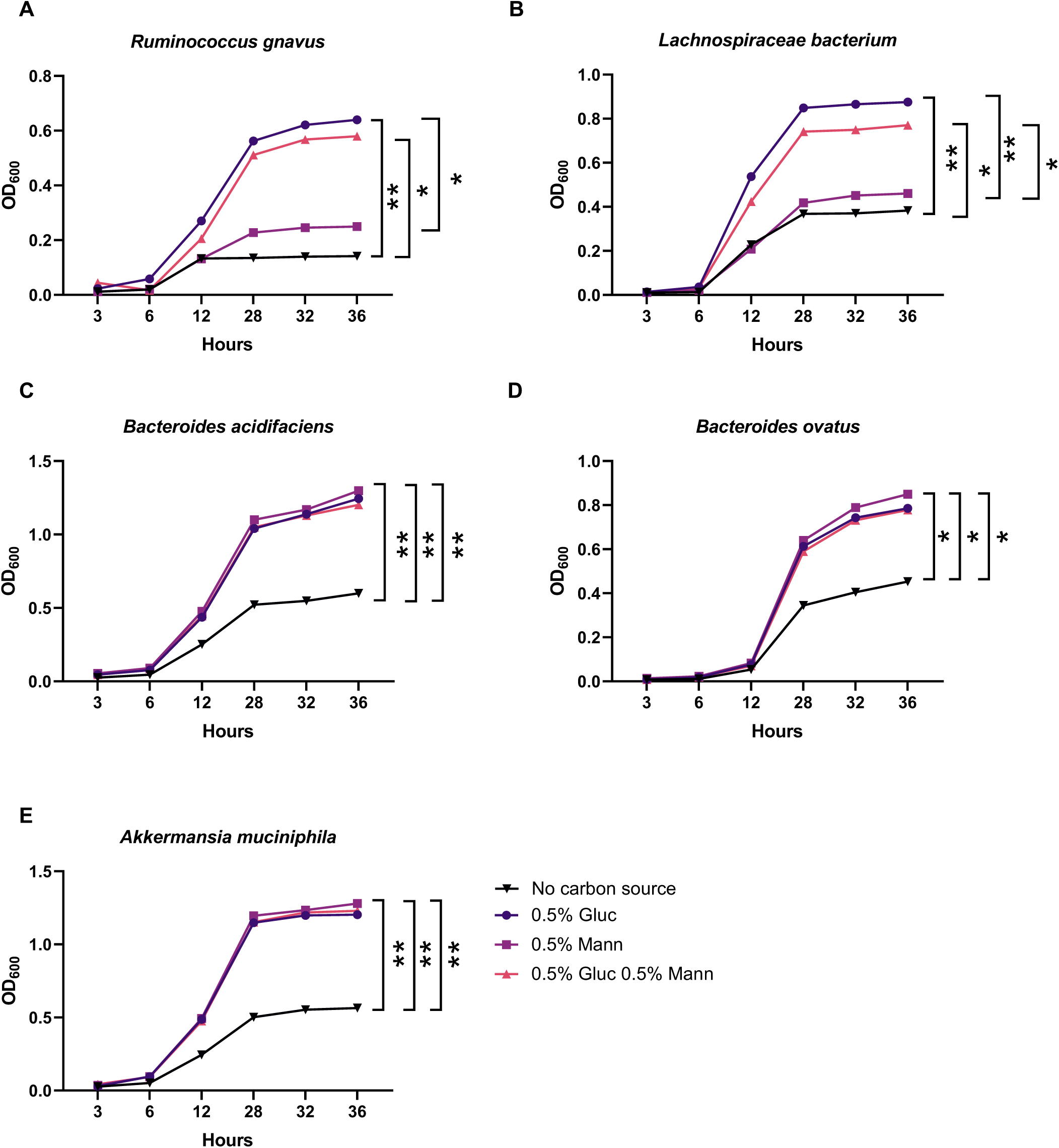
D-mannose suppresses the growth of *Firmicutes* in culture. A. Representative growth curve of *Ruminococcus gnavus* cultured in the heart infusion media without sugar (no carbon source), with 0.5% D-glucose or 0.5% D-mannose, measured during different intervals. B. Representative growth curve of *Lachnospiraceae bacterium* cultured in the heart infusion media without sugar (no carbon source), with 0.5% D-glucose or 0.5% D-mannose, measured during different intervals. C. Representative growth curve of *Bacteroides acidifaciens* cultured in the heart infusion media without sugar (no carbon source), with 0.5% D-glucose or 0.5% D-mannose, measured during different intervals. D. Representative growth curve of *Bacteroides ovatus* cultured in the heart infusion media without sugar (no carbon source), with 0.5% D-glucose or 0.5% D-mannose, measured during different intervals. E. Representative growth curve of *Akkermansia muciniphila* cultured in the heart infusion media without sugar (no carbon source), with 0.5% D-glucose or 0.5% D-mannose, measured during different intervals. Data are representative of one out of at least 3 independent experiments.

## Discussion

Obesity, type 2 diabetes, and metabolic syndrome are major health problems worldwide that pose financial and physical burdens to both affected individuals and society as a whole (1). Thus, understanding the causes of these disorders and developing new approaches to prevention and treatment takes high priority. Achieving these goals has been difficult, however, due to the complex interactions of the genetic and environmental factors driving these diseases, including host genetics, immune responses, diet, and the role of the gut microbiota. Here, we explored a novel way to treat obesity and type 2-like diabetes in mice caused by antibiotic treatment during the neonatal age. We revealed that D-mannose feeding, an epimer of glucose that shows therapeutic effects on experimental type I diabetes and airway inflammation in mice (9), suppresses the development of obesity and ameliorates metabolic abnormalities such as insulin resistance and glucose intolerance in NeoATB mice. Mechanistically, D-mannose restores the defective Tregs, consequently inhibiting the abnormally increased IFN-γ-type inflammatory T cells and proinflammatory M1-like macrophages. Importantly, we have discovered that D-mannose corrects the obesity-associated dysbiosis of the gut microbiota, specifically by adjusting the *Firmicutes*: *Bacteroidetes ratio* in obese NeoATB mice.

Most significantly, our findings indicate that D-mannose selectively suppresses the growth of *Firmicutes*, increasing the abundance of the phyla *Bacteroidetes* and *Verrucomicrobia* in both NeoATB and high-fat diet-obese mice.

### D-Mannose feeding suppresses obesity and improves metabolism in obese mice

Several conclusions can be drawn from the current study. D-mannose decreases body weight and reduces obesity in NeoATB mice without significantly affecting the overall weight of normal mice. D-Mannose-induced weight loss and obesity suppression was not due to reduced food intake in mice. The generality of D-mannose in suppressing obesity was confirmed in another obesity model of HFD diet feeding in mice. This conclusion is supported by several lines of cellular and molecular evidence. For example, obesity is accompanied by adipose tissue hypoxia, leading to an increase in the level of HIF1α (34, 35). The increase in HIF1α is linked to adipose tissue inflammation and insulin resistance (18). In the fat of obese NeoATB mice, we found increased HIF1α and reduced spare respiratory capacity of adipocytes (as a consequence of hypoxia), which may drive the increase in inflammation of adipose tissue, as determined by increased inflammatory cytokines including IFNγ, TNF, IL1β, IL-6, and IL-12, and highly accumulated Th1 type cells and M1-like macrophage. D-mannose treatment ameliorated hypoxia and reduced inflammation in adipose tissue of obese NeoATB mice.

### D-Mannose suppresses fat inflammation by increasing ST2^+^ Tregs

The question arises as to the mechanism by which D-mannose reduces fat inflammation and hypoxia? One possible mechanism is to increase Treg cells in fat tissue induced by D-mannose treatment, because the role of adipose Tregs, especially ST2^+^ Treg cells, in suppressing fat inflammation is well established. Loss-of-function and gain-of-function experiments have revealed that Treg cells control the inflammatory state of adipose tissue, thereby influencing insulin resistance (19). Here, we demonstrated that D-mannose treatment reversed the defect in the number of adipose ST2^+^Treg cells in the VAT of NeoATB mice, thereby abolishing inflammation in the adipose tissue. The indispensable role of ST2^+^ Treg cells in D-mannose-mediated suppression of inflammation and obesity in obese NeoATB mice was established when ST2^+^ Treg cells were depleted, which nullified the effect of D-mannose treatment. We sufficiently depleted ST2^+^Foxp3^+^ Treg cells in the adipose tissue with anti-CD25 antibody treatment, because adipose Treg cells co-express CD25 and ST2, and the antibody eliminated the majority of ST2^+^ Tregs. We did not use *Il1rl1^-/-^* (ST2 KO) mice in the D-mannose treatment experiment, as the ST2 KO mice have intrinsically reduced frequency and total number of adipose Treg cells and show increased weight of adipose tissue and impaired glucose tolerance, even on a normal diet (20) (36). The intrinsic feature of ST2 KO mice prevented us from testing the effect of D-mannose on the induction of adipose ST2^+^ Treg cells.

D-Mannose-mediated increase in adipose ST2^+^Treg cells in obese NeoATB mice can be attributed to multiple mechanisms. First, D-mannose drives more ST2^+^ Treg cell expansion in the fat tissue. Increase expression of ST2 on adipose Treg cells mediated by D-mannose can enhance their proliferation in response to IL-33, although we did not observe a significant difference in adipose IL-33 protein after D-mannose treatment. In addition, D-mannose may also promote the generation of more Treg cells in adipose tissue. D-mannose treatment increased TGF-β receptor I expression in fat CD4^+^CD25^-^ T cells, which then enhanced the sensitivity of these T cells to convert into Tregs (21). Indeed, we have shown that adipose Tregs could be converted from naïve CD4^+^ T cells in the adipose tissue (Zanvit et al, in revision). Furthermore, D-mannose-mediated suppression of inflammation in the adipose tissue may also be attributed to an increase in ST2^+^ ILC2s and a decrease in macrophages, which may not necessarily be dependent on Tregs. Although it remains elusive, the up-regulatory effects on Treg and ILC2 cells in adipose tissue could also be mediated by the regulatory function of D-mannose on gut microbiota. Furthermore, D-mannose treatment reduced macrophage infiltration in the adipose tissue of obese mice, which was Treg-independent. We also observed decreased expression of *II1b* and *Hif1a* mRNAs in adipose tissue of D-mannose-treated NeoATB mice. This data is consistent with recent findings that D-mannose can suppress expression of IL-1β through downregulation of HIF1α in macrophages (37).

### D-Mannose corrected microbiota dysbiosis in the gut of obese NeoATB mice

D-Mannose treatment corrected the dysbiosis of gut microbiota in obese NeoATB mice. The gut microbiota plays a fundamental role in the regulation of the immune response and metabolism. An altered (decreased) ratio of *Bacteroidetes*:*Firmicutes* was reported in both diet-induced and genetically obese mouse models and humans (25, 38). Consistent with these reports, we observed a reduced ratio of *Bacteroidetes* to *Firmicutes* in obese NeoATB mice, as well as in the HFD-induced obesity model. D-Mannose treatment significantly suppressed the abundance of phylum *Firmicutes* and increased the abundance of phylum *Verrucomicrobia* (in particular *Akkermansia muciniphila*) and *Bacteroidetes* (39). Family *Ruminococcaceae* represents one of the most abundant families of phylum *Firmicutes* that was previously linked with obesity (40). In both NeoATB and HFD-fed obese mice, D-mannose treatment significantly reduced the abundance of the family *Ruminococcaceae*. Our findings also provide the underlying mechanism for a recently reported positive effect of D-mannose on HFD-induced obesity, in which the authors show a preventive but not therapeutic effect of D-mannose on the reduction of obesity in an HFD-induced obesity model (39).

In obese NeoATB mice, based on predicted microbiota function (PICRUSt, level 3), we identified significantly increased pathways such as ABC transporters, bacterial chemotaxis, flagellar assembly or phosphotransferase system (PTS), that offer a potential functional prediction of the microbiota as previously identified in obese Chinese children and adolescents (3-18 years of age) (41). Significantly, D-mannose treatment normalized all aforementioned pathways to the frequency observed in the control group. Obese patients following weight-loss intervention showed significantly increased prediction of the oxidative phosphorylation pathway compared to significantly increased flagellar assembly and bacteria chemotaxis pathways detected in weight-loss intervention unsuccessful patients with metabolic syndrome (42). Consistent with this observation in humans, D-mannose-treated NeoATB mice had significantly increased prediction of the oxidative phosphorylation pathway. Moreover, D-mannose treatment of obese NeoATB mice corrected other metabolic pathways that were previously associated with obesity, such as the pentose phosphate pathway, biotin metabolism pathway, folate biosynthesis pathway, or the tryptophan metabolism pathway (43–46). Nevertheless, the effect of D-mannose on the regulation of metabolic pathways should be investigated in more detail in future studies.

### D-Mannose suppresses Propionate in obese NeoATB mice

Obesity was not only linked to the composition of the microbiota but also to the production of short-chain fatty acids derived from the gut microbiota. The phylum *Firmicutes* represents one of the most important phyla that produces large amounts of SCFAs, which are increased in the obese but not in the lean subjects. Interestingly, the proportion of individual SCFAs changed in favor of propionate in overweight and obese people (31). Consistent with human data, obese NeoATB mice had significantly increased fecal concentrations of propionate compared to control mice, although other SCFAs were also elevated. This increase in propionate was significantly reduced by D-mannose treatment in the NeoATB mice, which was likely attributable to a reduced abundance of the phylum *Firmicutes.* The functional significance of D-mannose-mediated decrease in propionate was further illuminated by the reduction of FABP4, which is known to increase glucagon and impair insulin action in mice and humans, and is thus positively linked to obesity (32). Interestingly, we have also observed that treating young NeoATB mice with D-mannose prevents the development of obesity in adulthood. This effect was canceled if mice were treated with the combination of D-mannose and Propionate.

Our data further revealed a decrease in fecal concentration of Butyrate in NeoATB mice treated with D-mannose. The role of Butyrate in obesity is complex and multifaceted. While it has the potential to improve obesity, substantial evidence also suggests it may contribute to the condition. Obese individuals commonly exhibit a higher ratio of Firmicutes:Bacteroidetes in their gut microbiota, with Firmicutes being key producers of Butyrate. Interestingly, research by Schwiertz et al. (2010) found increased Butyrate levels in obese patients, linked to higher *Bacteroidetes* levels. Additionally, Butyrate can stimulate lipid synthesis through the β-hydroxy-β-methylglutaryl-CoA pathway, which may further promote obesity (Birt DF, 2013). These findings highlight the need for further investigation into the dual role of Butyrate in relation to obesity.

### D-Mannose specifically suppresses the growth of *Firmicutes*

The inverse correlation between the altered ratio between *Firmicutes* and *Bacteroidetes* in the feces of D-mannose-treated obese mice revealed that D-mannose directly and specifically inhibits the growth of bacteria that belong to the phylum *Firmicutes*. In a series of *in vitro* experiments, we cultured representative bacteria selected from the phyla Firmicutes, Bacteroidetes, and Verrucomicrobia. We found that D-mannose selectively inhibits the growth of Firmicutes members, specifically Ruminococcus gnavus and a *Lachnospiraceae bacterium*. This effect is specific to D-mannose, as the same concentrations of glucose in the culture promote the growth of *Ruminococcus gnavus* and *Lachnospiraceae bacterium*. On the other hand, D-mannose does not suppress the growth of *Verrucomicrobia* member *Akkermansia muciniphila* and the *Bacteroidetes* members *Bacteroides acidifaciens* and *Bacteroides ovatus*. This finding is significant because it not only provides a mechanism for the decreased ratio of phylum *Firmicutes* to *Bacteroidetes* or *Firmicutes* to *Verrucomicrobia* in the gut of D-mannose-treated NeoATB mice but also reveals previously unrecognized selective effects of D-mannose on the gut microbiota. This could have potential translational implications for clinicians to manipulate the gut microbiota for the development of immunotherapy for human obesity and type 2 diabetes with a single sugar. Indeed, several reports have identified positive effects of *Akkermansia muciniphila* on the reduction of obesity since its fecal abundance was positively correlated with leanness in mice and humans (47). A very recent study demonstrated the potential application of *Akkermansia muciniphila* as an effective probiotic treatment of human obesity (48). Thus, we believe that D-mannose-mediated improvement of obesity in both NeoATB and HFD-fed mice is an exciting pre-clinical model for developing related therapy for human obesity by manipulating gut microbiota.

## Conclusion

Collectively, our data reveal a significant effect of D-mannose on the treatment of obesity. This effect was achieved through modification of the gut microbiota and related SCFA, as well as increased adipose Treg cells, which, in turn, resulted in a reduction of adipose inflammation and improved metabolism. We believe that D-mannose treatment may have therapeutic implications for obesity in human patients. Clinical trials will be vital, thereby enabling the development of new and effective approaches for the treatment of obesity and insulin resistance.

### Experimental Procedures

#### Mice

C57BL/6J male mice were used in all experiments. Mice were bred under specific pathogen-free conditions in the animal facility of the National Institute of Dental and Craniofacial Research. All animal studies were performed according to National Institutes of Health guidelines for the use and care of live animals and were approved by the Animal Care & Use Committee (ACUC) of the National Institute of Dental and Craniofacial Research. We have complied with all relevant ethical regulations for animal testing and research.

#### Mice treatments

Neonatal antibiotic treatment was performed using vancomycin hydrochloride (500 mg/L, Hospira) and polymyxin B (1,000,000 U/L X-gen). Neonates were treated in intervals of 3 weeks, starting immediately after birth. After three weeks, the mice were weaned and housed in a standard environment with regular chow and water. After the onset of obesity, mice were treated with either water or a 20% D-mannose solution in water. In all experiments, only male mice were used. For the diet-induced obesity (DIO) model, 18-week-old male DIO C57BL/6NTac mice (Taconic Model #DIO-B6-M) were supplemented with a high-fat diet (60% kcal% fat) obtained from Research Diets (catalog# D12492) and supplemented with either normal drinking water or 20% D-mannose in water. The weight of the mice was monitored during the experiment. Tissues were harvested at the end of the experiments for histopathological and immunological analyses.

#### ELISA

The levels of IL-33 in tissue supernatants were determined using a sandwich Enzyme-Linked Immunosorbent Assay (ELISA) kit, as per the manufacturer’s instructions (Invitrogen by Thermo Fisher Scientific). Briefly, a standard 96-well plate (Nunc-Immuno MaxiSorp Uncoated Plates) was coated with the appropriate capture antibody and incubated at 4°C overnight. After washing, the non-specific binding was blocked by adding assay diluent. Standards and tissue supernatants were dispensed into the wells and after 1.5 h of incubation, the detection antibody was added to each well for 1 h. After washing avidin-HRP enzyme conjugate was added into each well. Thirty min later, the plate was washed and incubated with TMB substrate solution at room temperature in the dark until an adequate signal developed. The reaction was stopped by 2 N H_2_SO_4_ and the optical density was determined using the microplate reader Spectramax Plus (Molecular Devices, San Jose, CA). The concentration of IL-33 in each sample was interpolated from standard curves and expressed as pg of cytokine per mL. To determine levels of total and active TGFβ1, TGFβ1 E_max_ ImmunoAssay System was used (Promega) following manufacturer’s instructions.

#### Bacterial cultures

The bacterial strains Bacteroides acidifaciens (DSM 15896), Bacteroides ovatus (ATCC 8483), Akkermansia muciniphila (ATCC BAA835), Ruminococcus gnavus (ATCC 29149), and Lachnospiraceae bacterium (DSM 24404) were cultured following ATCC or DSMZ instructions. *Bacteroides acidifaciens*, *Bacteroides ovatus*, *Ruminococcus gnavus*, and *Lachnospiraceae bacterium* were cultured in chopped meat medium (Anaerobe systems). *Akkermansia muciniphila* was cultured in Brain Heart Infusion medium (BD Bacto Brain Heart Infusion, Thermo Fisher Scientific). After culture in chopped meat media, *Bacteroides acidifaciens*, *Bacteroides ovatus*, *Ruminococcus gnavus*, and *Lachnospiraceae bacterium* were further cultured in brain heart infusion. Heart infusion media (BD Bacto, Heart infusion broth, Thermo Fisher Scientific) (containing no dextrose) were used as a minimal medium, without any sugar (no carbon source) or supplemented with 0.5% D-glucose or 0.5% D-mannose. Bacterial growth was determined by measuring OD at 600nm wavelength in 1mL of bacterial suspension at different time points.

#### Glucose tolerance test and Insulin tolerance test

Mice underwent metabolic studies beginning at about 7 days before the end of experiments, when the glucose tolerance test (GTT) was performed. Mice were fasted overnight (15-16h) period, weighed, and then blood glucose concentrations were measured at 0, 15, 30, 60, 90, and 120 min after intraperitoneal glucose administration (2 g/kg). The insulin tolerance test (ITT) was performed ∼4 days after the GTT. For ITT, mice were fasted for 4-6 hours and injected intraperitoneally with insulin (Humulin R, Lilly; 0.75 U/kg). Glucose concentration was determined at 0, 20, 40, 60, and 80 min afterward.

#### Preparation and flow cytometry of adipose tissue samples

Mice were killed using CO_2_. Epididymal adipose tissue was removed and placed into Dulbecco’s modified Eagle’s medium (DMEM), by scissors, mechanically disaggregation of the tissue, and incubated with collagenase type II (1 mg/mL; Worthington Biochemical Corporation) at 37°C for 45 min. Single-cell suspensions were obtained by mincing the tissues through 70-μm cell strainers and further washed with DMEM. For flow cytometry involving surface staining only, cell preparations were subjected to red blood cell lysis by incubating with 1 mL ammonium-chloride-potassium lysing (ACK) buffer for 2-3 min on ice, washed, and finally resuspended in DMEM. Live/dead staining was performed before surface staining using the Zombie Yellow Fixable Viability Kit (BioLegend) for 10 minutes at room temperature. Surface staining with anti-mouse-specific antibodies was performed for 20 minutes at 4°C. Following standard surface staining, intracellular staining of transcription factors was performed using the Foxp3/Transcription Factor Staining Buffer Set (Thermo Fisher Scientific), followed by staining with anti-mouse-specific antibodies (such as anti-mouse Foxp3 or anti-mouse Gata3). For intracellular cytokine staining, cells were stimulated for 4 h at 37°C with PMA (phorbol 12-myristate 13-acetate; 5 ng/ml), Ionomycin (1 μg/ml), and Golgi-Plug (1:1,000 dilution; BD Pharmingen), followed by BD Cytofix/Cytoperm permeabilization buffer according to the manufacturer’s instructions (BD Biosciences). Cells were further stained with specific antibodies to determine the expression of cytokines. A list of all antibodies used for flow cytometry is shown as Supplemental Table 1.

#### PCR quantification

RNA from adipose tissue was extracted using a protocol described previously (49). RNA from *ex-vivo* sorted adipose T cells or macrophages was extracted using RNeasy Plus Micro Kit (Qiagen). cDNA was synthesized using a High-Capacity cDNA Reverse Transcription Kit (Applied Biosystems by Thermo Fisher Scientific). In samples with low concentrations of RNA/cDNA (mostly from sorted cells), pre-amplification was performed using TaqMan PreAmp Master Mix (Applied Biosystems by Thermo Fisher Scientific). qPCR was done using TaqMan assays following the manufacturer’s instructions. All TaqMan assays are listed in the Supplemental Table. 2. Total transcript values were normalized to those from mouse *Hprt* mRNA. Results were calculated using the comparative ΛιΛιCT method.

#### Western blot

Tissue lysates from the adipose tissue were prepared as described previously (50). Total protein concentration was determined using a Bradford Protein Assay (Bio-Rad). Protein samples were separated either on 4-12% Bis-Tris or 4-20% Tris-Glycine gels (Thermo Fisher Scientific) and transferred to 0.45 µm nitrocellulose membranes (Thermo Fisher Scientific). The membranes were soaked in blocking buffer for 1 h at room temperature and subsequently incubated with the appropriate primary antibody overnight at 4°C. The next day, the membranes were washed and incubated for 1 h at room temperature with HRP-conjugated secondary antibodies (Santa Cruz Biotechnology). Immunoreactivity was detected using Super Signal West Pico or Dura Chemiluminescent Substrate (Thermo Fisher Scientific), followed by stripping the membranes with Reblot Plus Strong Solution (Merck Millipore) and incubated with α-tubulin antibody (Sigma-Aldrich) as a control. The list of the antibodies used for the Western blot is in Supplemental Table 3.

#### Seahorse Assay

Oxygen consumption rate (OCR) was measured using a 96 well Extracellular Flux Analyzer (Seahorse bioscience), which was described previously (51). 5-8 mg of epididymal fat were attached onto the wells of a Seahorse X96 spheroid microplate coated with Poly-L-Lysine (50 μg/mL for 20 min at room temperature). The plate was preincubated at 37°C for 45 min, in the absence of CO_2_, in RPMI medium pH 7.4 supplemented with 10 mM glucose, 1 mM pyruvate, and 2 mM L-glutamine. OCR was measured under basal condition (0-60 min), after Oligomycin stimulation (60-100 min, 8 µM), CCCP stimulation (100-140 min, 8 µM) and Rotenone/Antimycin-A stimulation (140-200 min, 3 and 12 µM respectively; all the reagents were from Sigma). Results were collected with Wave software 2.4 (Agilent technologies). All reagents used for the Seahorse assay are listed in Supplemental Table 4.

#### 16S sequencing

The determination of the total bacterial load was performed via qPCR using 16S rRNA gene primers, and *Streptococcus gordonii* (ATCC 35105) genomic DNA was used as a standard. The V1-V2 or V3-V4 region of the 16S rRNA gene was amplified from fecal gDNA. DNA was extracted using QIAamp DNA Stool Mini Kit (Qiagen) according to the manufacturer’s instructions. Sequencing was performed on an Illumina MiSeq using the V3 reagent kit (600 cycles). The sequencing was performed with >10% PhiX spike-in. The Nephele (52) (https://nephele.niaid.nih.gov) was used to demultiplex and cluster sequences into 99% identity OUT’s (QIIME pipeline) and taxonomic annotation assigned based on the open Greengene database. Abundance, diversity plots and LEfSe (LDA) were calculated using Microbiome analyst project (53) (https://www.microbiomeanalyst.ca). All plots showing microbiota abundance and diversity were generated in R (54).

#### Functional prediction of gut microbiota

The prediction of the functional genes in the gut microbiota was done using the PICRUSt (27). The taxonomy assignment (Greengene database with a 97% similarity) as well as PICRUSt workflow was done using the Nephele platform (52) (https://nephele.niaid.nih.gov) and final metagenome functional prediction from the Kyoto Encyclopedia of Genes and Genomes (KEGG) database at hierarchy level 3 pathways were obtained. Statistical Analysis of Metagenomic Profiles (STAMP) software v2.1.3 was utilized to analyze the PICRUSt-predicted metagenomes (28). Significant differences in the functional genes between the groups were identified using the One-way ANOVA test, followed by FDR (Benjamini Hochberg) posttest for multiple test correction.

#### Fecal Microbiota Transfer

For the fecal microbiota transplant (FMT) experiment, we first collected fecal samples from NeoATB mice or Mannose-treated NeoATB mice, both aged 24 to 26 months. Samples were stored at -80 °C until use. On the day of the FMT, the feces were thawed, and 200 mg of the fecal material was diluted in 2 mL of phosphate-buffered saline (PBS). The mixture was then homogenized and thoroughly mixed. Next, the fecal suspension was filtered, and centrifugation was performed at 600g for 5 minutes to pellet any undissolved solid matter. The supernatant was then garaged to the recipient mice at a dosage of 10 μL per gram of body weight, administered every other day for a total of three doses.

#### Chromatographic and mass detection analysis

Short-chain fatty acid (SCFA) acetate, propionate, and butyrate chromatography was carried out on a capillary column (30 m × 0.250 mm, 0.25 μm; Agilent Technologies) with a 34.5 min run time. SCFA were separated at 2.61 min retention time, m/z 60 (qualifier ions m/z 43, 29) for acetate; 2.65 min retention time, m/z 74 (qualifier ions m/z 57, 45) for propionate; 2.71 min retention time, m/z 60 (qualifier ions m/z 88, 73) for butyrate, protonated form on SIM mode of 20–550 m/z. Analyses was performed with an Agilent 6890N gas chromatograph coupled to an Agilent 5973 mass-selective detector (MSD) with the following chromatographic conditions: Initial temperature 100°C for 0.50 min, increasing to 180°C at 8°C/min for 21 min and finally increased 200°C at 20°C/min for 2 min. The front inlet temperature was 240°C, operating with a split-less operation mode. MSD ion source and interface temperature were 230°C and MS Quad at 150°C. The MSD operated in EI mode at 70 Ev and 1306 relative EMV volts. Carrier gas was He (1.0 ml/min). GC-MS data were acquired and processed using Agilent MassHunter WorkStation Software.

#### Sample Preparation for GC-MS analysis

Fecal pellets were placed into 0.3 ml of deionized water and followed by homogenization using a Precellys homogenizer (Bertin Instruments), utilizing 1.0 mm zirconia/silica beads for 3 X 40 sec at 6500 rpm. The pH of the suspension was adjusted to 2.2 ± 0.05 by adding 5M HCl at room temperature for 20 min with sonication. The samples were centrifuged at 20000g for 15 min at 4°C. Furthermore, 0.2 ml of supernatant was taken into a GC-MS Vial and 2 μl was injected into the GC-MS using an autosampler. Samples were analyzed as above in GC-MS. Final concentrations were normalized to the weight of fecal pellets.

#### Histology

Adipose or liver tissues were collected and fixed in 10% Neutral Buffered Formalin. All tissues were processed at the Histoserv company (https://www.histoservinc.com). Two sections per tissue were stained with hematoxylin and eosin for analysis. Images were acquired with the Leica Aperio Scan Scope instrument and analyzed using Aperio Image Scope Software. Adipocyte size was determined using AdipoCount Software (55). Liver steatosis histology scoring was done blindly by two independent investigators and scored based on the scoring described previously (56).

#### Statistical analysis

All statistical analyses were performed using GraphPad Prism 8 software. All data are presented as means ± S.E.M. Statistical significance, as indicated by asterisks, was determined by an unpaired t-test (two-tailed) or by one- or two-way ANOVA. p < 0.05 was considered significant (*, p < 0.05; **, p < 0.01; ***, p < 0.001; ****, p < 0.0001).

## Acknowledgments

This research was supported by the Intramural Research Programs of the National Institute of Dental and Craniofacial Research (NIDCR) and NIAID, National Institutes of Health (NIH), USA.

We want to thank the Combined Technical Research Core and the Veterinary Resources Core at NIDCR for their service and technical assistance. We want to thank Robert J Palmer for the help and suggestions with bacterial culture experiments.

## Author Contributions

P.Z. designed and did experiments, analyzed data, and drafted the manuscript. J.X., N.G., D.Z., M.P., T.G., D.P.P., A.C., W.J., and F.J.G. designed and/or performed experiments. Y.B. provided critical scientific input. W.J.C. conceived of and supervised the research, designed the experiments, and wrote the manuscript.

## SUPPLEMENTAL FIGURES AND TABLES

**Supplemental Figure 1.**
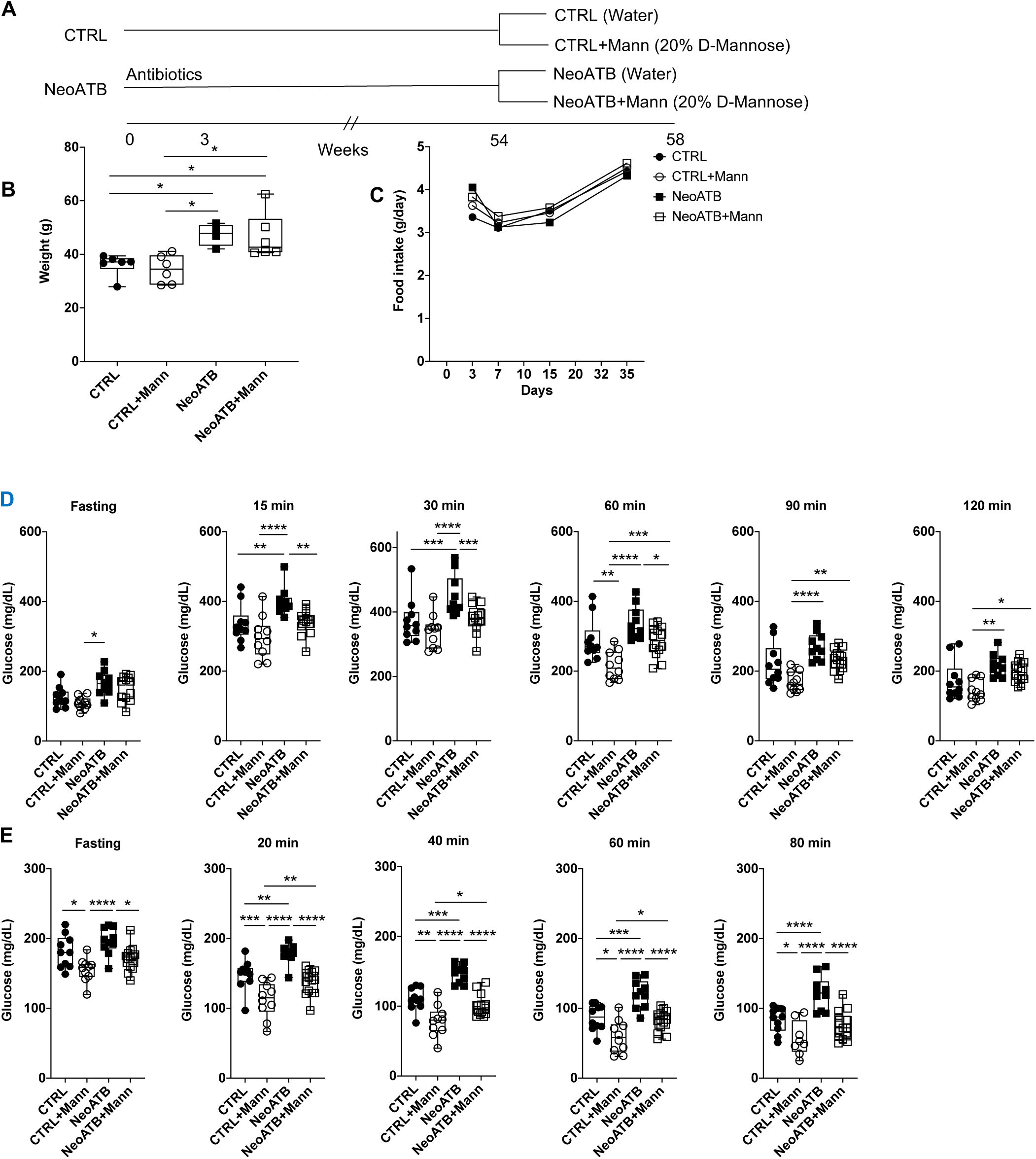
D-mannose improves metabolism of NeoATB mice. a) Immunization scheme. b) Weight of control and obese NeoATB mice at 54 weeks of age before D-mannose treatment. c) Food intake of Control, Control treated with D-mannose, NeoATB and NeoATB treated with D-mannose mice during interval of 35 days. d) Glucose tolerance test of Control, Control treated with D-mannose, NeoATB and NeoATB treated with D-mannose mice. e) Insulin tolerance test of Control, Control treated with D-mannose, NeoATB and NeoATB treated with D-mannose mice Data in d) and e) are pooled of 2 independent experiments. Each point in figures represents one individual animal. Statistical analysis was performed using one-way ANOVA (* p<0.05; ** p<0.01; *** p<0.001; **** p<0.0001).

**Supplemental Figure 2.**
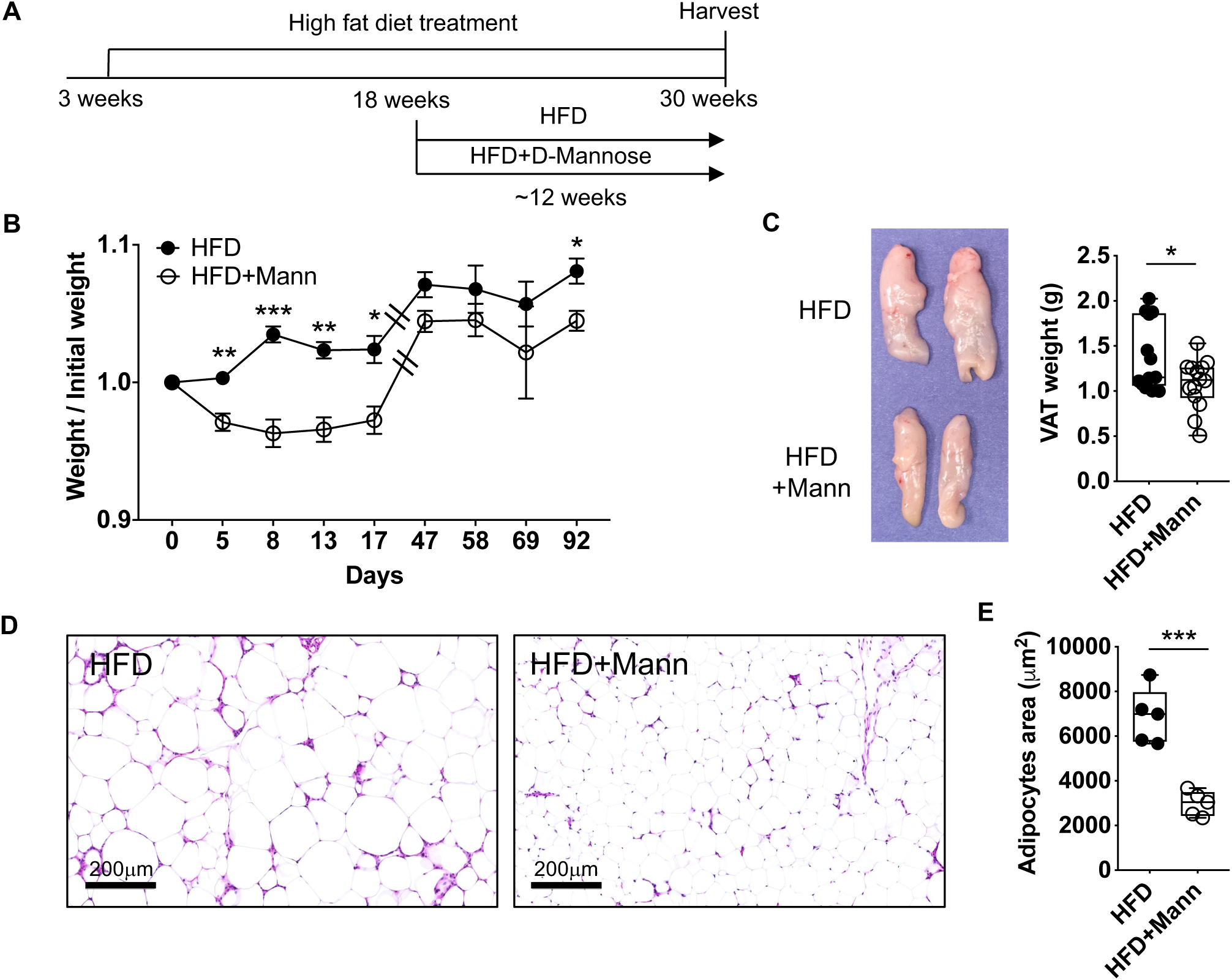
D-mannose ameliorates obesity in mice treated with a high-fat diet. A. Immunization scheme. B. Weight/Initial weight ratio is shown between HFD and HFD+D-Mannose treated mice. C. Representative photograph of visceral adipose tissue (VAT) of HFD or HFD D-mannose-treated mice. Summarizing data showing the weight of VAT between HFD and HFD D-mannose-treated mice. D. Representative H&E staining of VAT in HFD and HFD D-mannose-treated mice. E. Summarizing data showing the size of adipocyte area between HFD and HFD D-mannose-treated mice determined using AdipoCount software. Data are pooled from 2 independent experiments. Each point in the figures represents one individual animal. Statistical analysis was performed using a two-tailed *t-test* (* p<0.05; ** p<0.01; *** p<0.001).

**Supplemental Figure 3.**
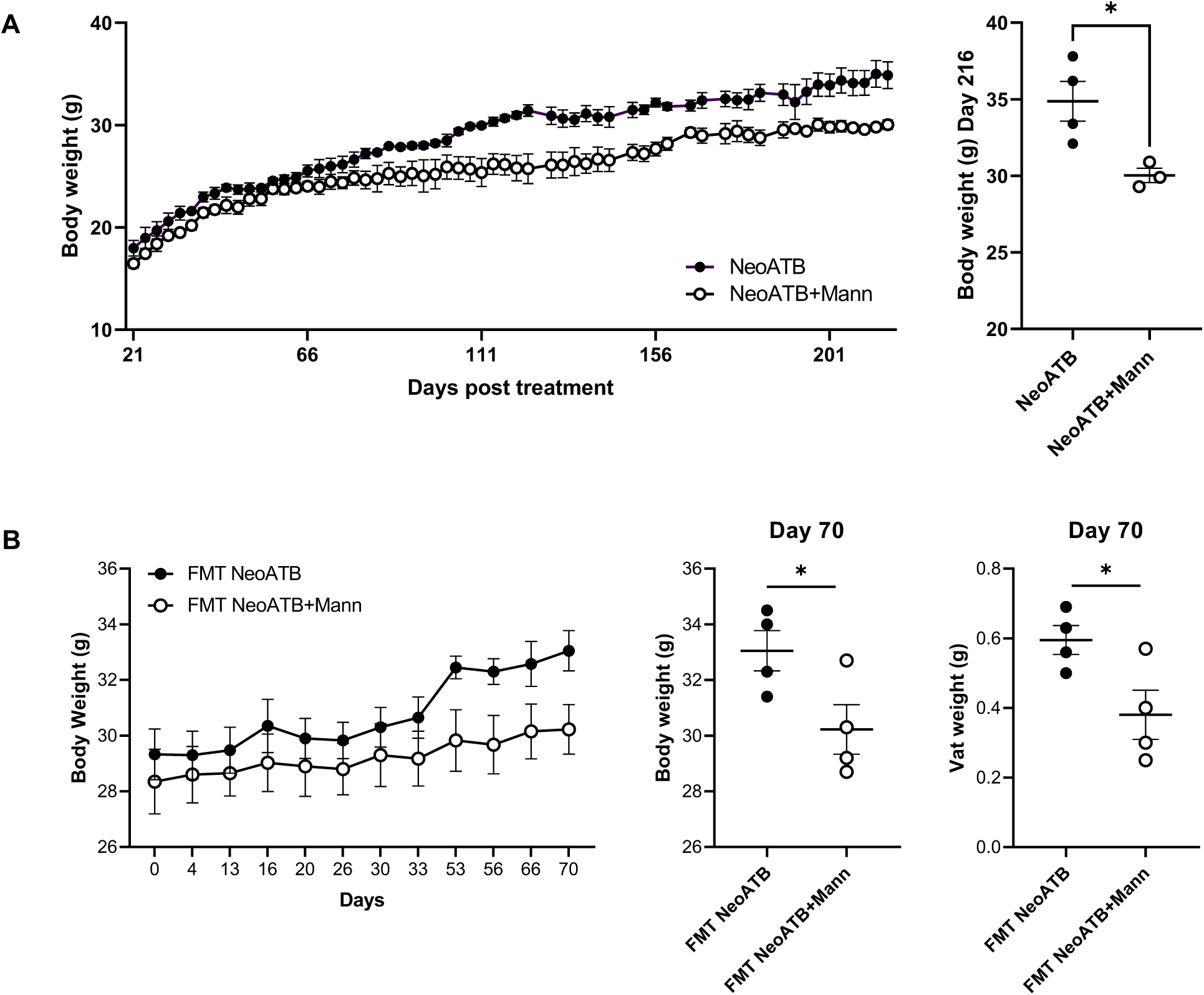
A. D-Mannose treatment can ameliorate the development of obesity in young NeoATB mice treated with antibiotics at a neonatal age. Mice were treated with vancomycin and polymyxin B during neonatal life (the first 3 weeks). After weaning, mice were separated into two groups: NeoATB (n=4) and NeoATB treated with 20% D-mannose (NeoATB+Mann; n=4) in drinking water. Body weight was monitored for 200+ days. B. FMT transfer of microbiota from NeoATB mice treated with D-mannose ameliorated obesity in adult NeoATB mice. For the fecal microbiota transplant (FMT) experiment, we first collected fecal samples from NeoATB mice or Mannose-treated NeoATB mice, both aged 24 to 26 months. These samples were stored at -80 °C until use. On the day of the FMT, the feces were thawed, and 200 mg of the fecal material was diluted in 2 mL of phosphate-buffered saline (PBS). The mixture was then homogenized and thoroughly mixed. Next, the fecal suspension was filtered, and centrifugation was performed at 600g for 5 minutes to pellet any undissolved solid matter. The supernatant was then gavaged to the recipient mice at a dosage of 10 μL per gram of body weight, administered every other day for a total of three doses. The data is representative of 1 experiment. Statistical analysis was performed using a two-tailed *t-test* or two-way ANOVA test (* p<0.05).

**Supplemental Figure 4.**
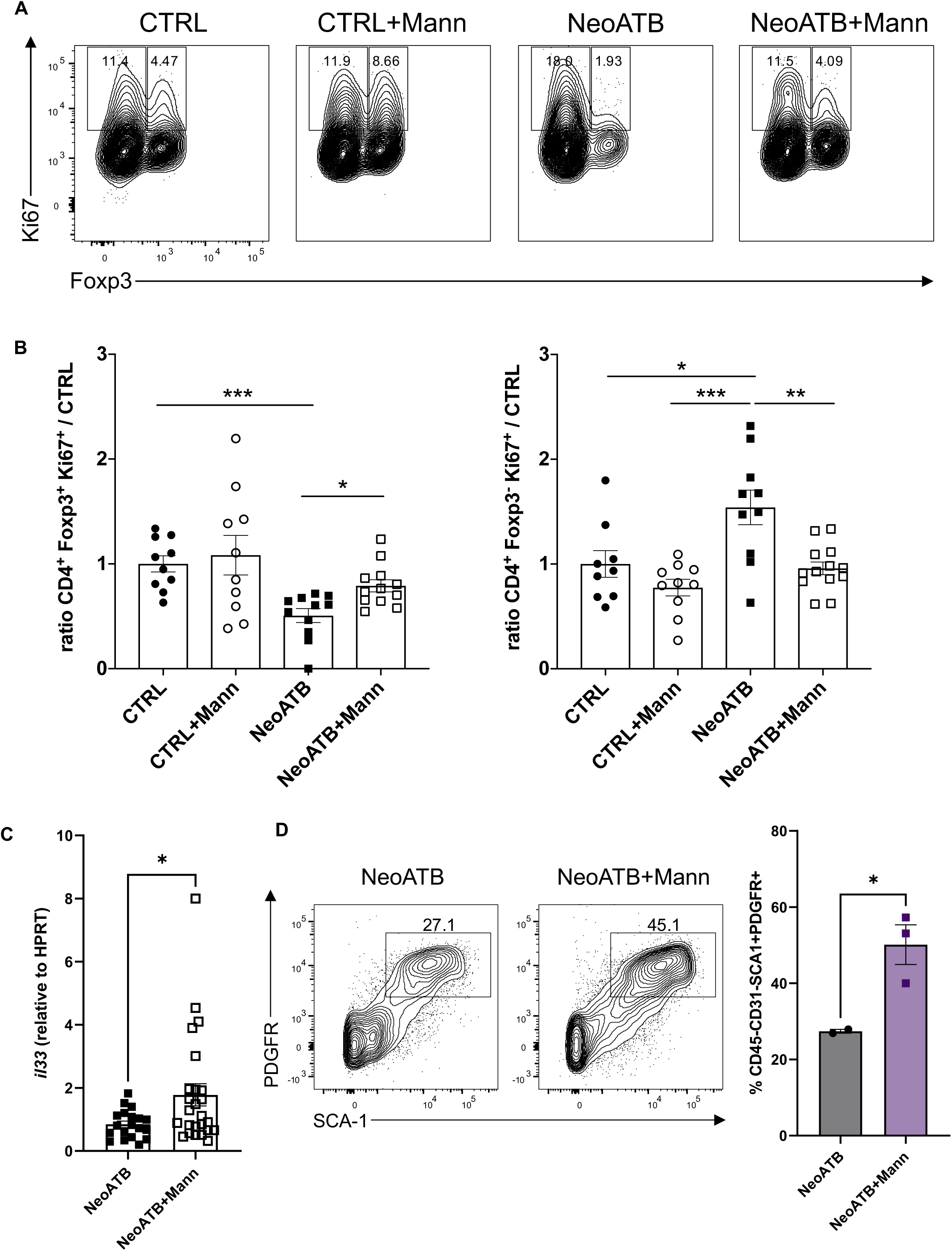
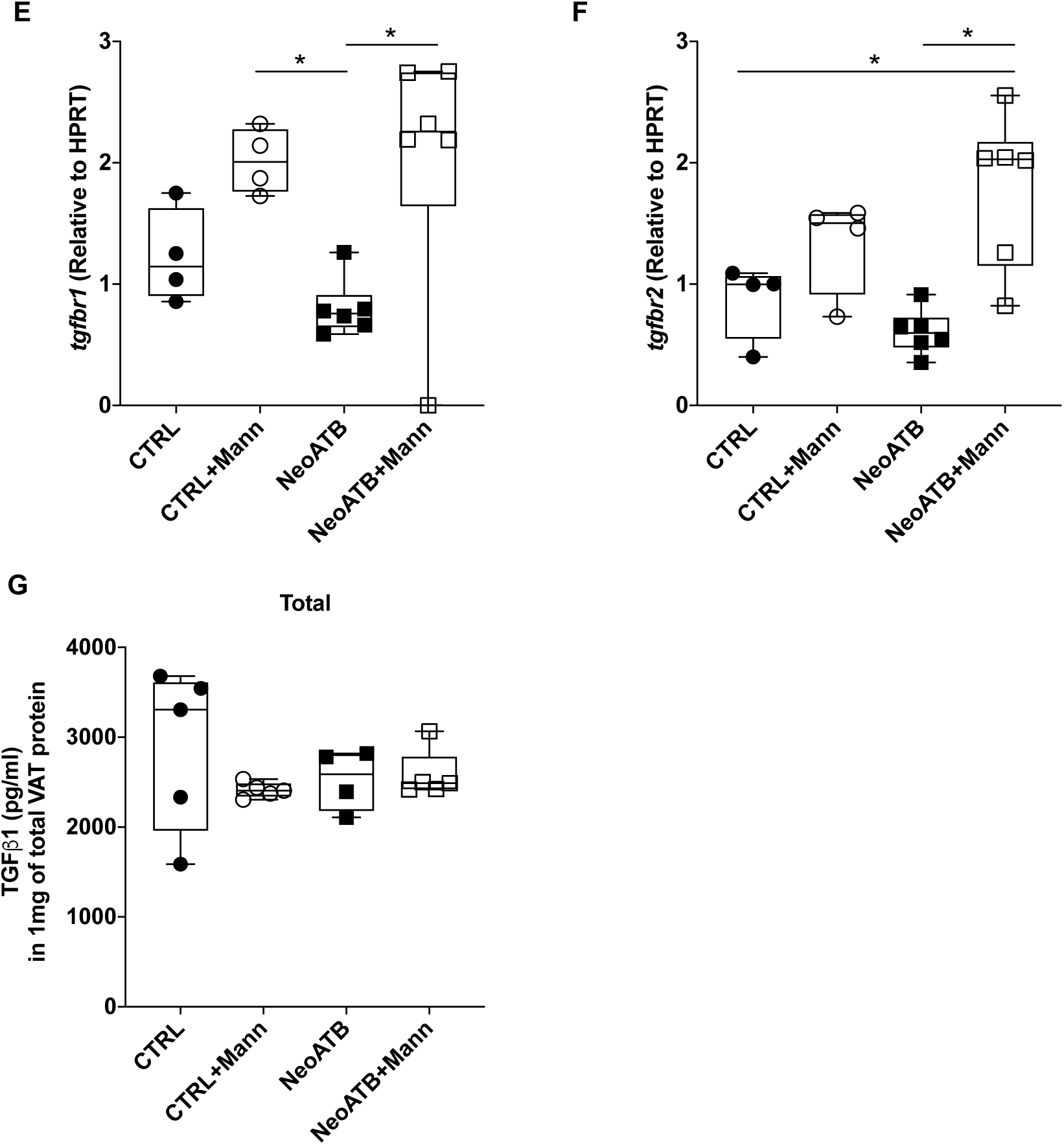
D-mannose treatment increases the proliferation of adipose Treg cells. A. Representative FACS plots showing the frequency of Ki67^+^Foxp3^+^ (Treg) or Ki67^+^Foxp^-^(Responder T) cells. B. Summarized data showing the ratio of CD4^+^Ki67^+^Foxp3^+^ to Control cells or the ratio of CD4^+^Ki67^+^Foxp3^-^ responder T cells. C. Summarized data showing gene expression of Interleukin-33 (*il33*) in the adipose tissue among different groups. D. Flow cytometric analysis of CD45-CD31-SCA1+PDGFR+ VAT mesenchymal cells, which represent the primary source of IL-33 in the VAT. E. Summarized data showing gene expression of *tgfbr1* in the adipose tissue among different groups. F. Summarized data showing gene expression of *tgfbr2* in the adipose tissue among different groups. G. Summarized data showing total concentration of total adipose TGFβ1 protein normalized to 1 mg of total fat protein, among different groups determined by ELISA. Data are pooled from 2-3 independent experiments (a, b, c, e, f, g) or 1 experiment (d). Each point in the figures represents one individual animal. Statistical analysis was performed using one-way ANOVA (* p<0.05; ** p<0.01; *** p<0.001).

**Supplemental Figure 5.**
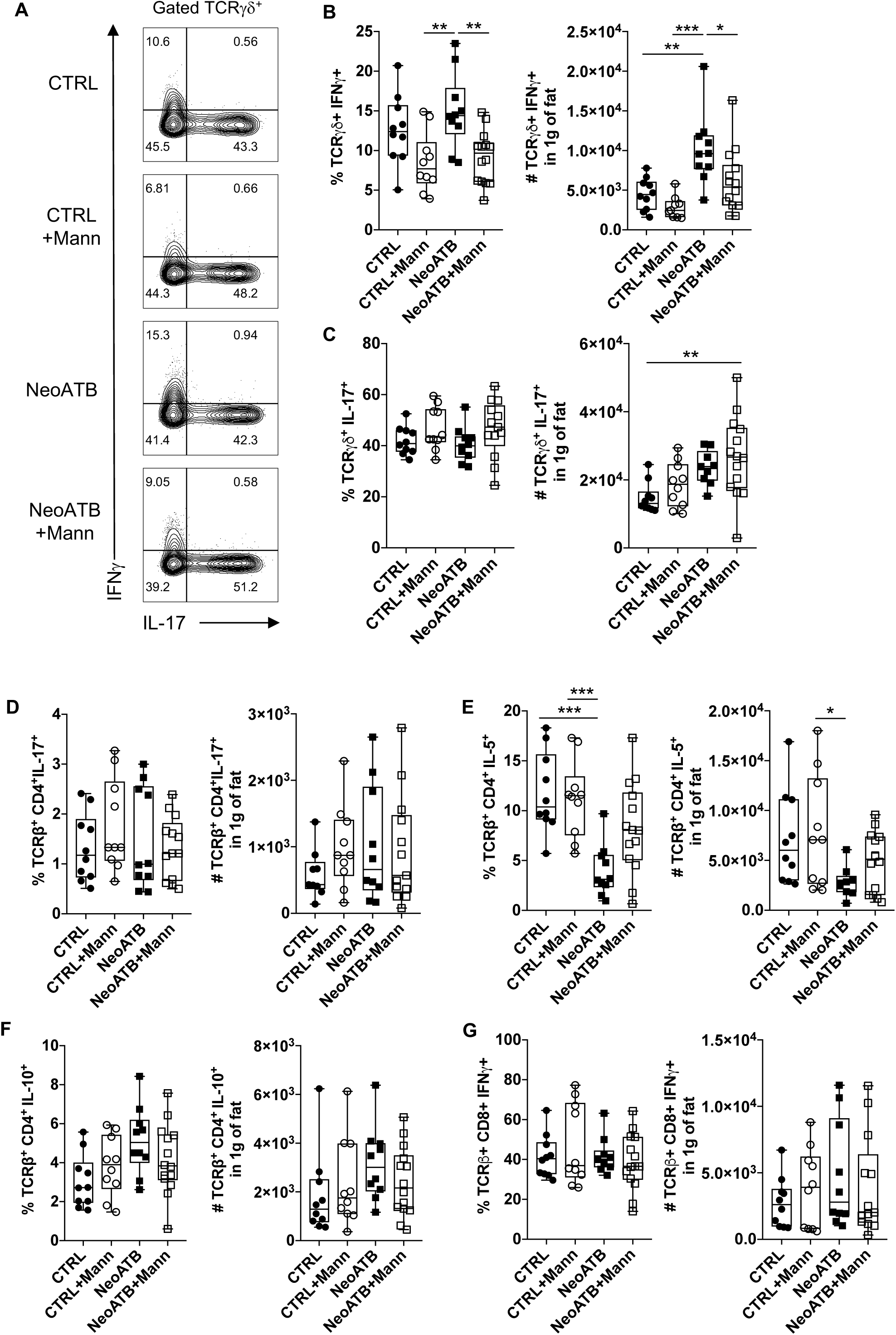
D-mannose decreases frequency and total number of adipose TCRγ8^+^IFNγ^+^ cells in NeoATB mice. A. Representative FACS plots showing frequency of adipose TCRγ8^+^IFNγ^+^ or TCRγ8^+^IL-17^+^ cells (Gated as: CD45^+^TCRγ8^+^) among different groups. B. Summarizing data showing frequency and total number of adipose TCRγ8^+^IFNγ^+^ cells among different groups. C. Summarizing data showing frequency and total number of adipose TCRγ8^+^IL-17^+^ cells among different groups. D. Summarizing data showing frequency and total number of adipose CD4^+^IL-17^+^ cells among different groups. E. Summarizing data showing frequency and total number of adipose CD4^+^IL-5^+^ cells among different groups. F. Summarizing data showing frequency and total number of adipose CD4^+^IL-10^+^ cells among different groups. G. Summarizing data showing frequency and total number of adipose CD8^+^IFNγ^+^ cells among different groups. Data are polled of 2-3 independent experiments. Each point in figures represents one individual animal. Statistical analysis was performed using one-way ANOVA (* p<0.05; ** p<0.01; *** p<0.001).

**Supplemental Figure 6.**
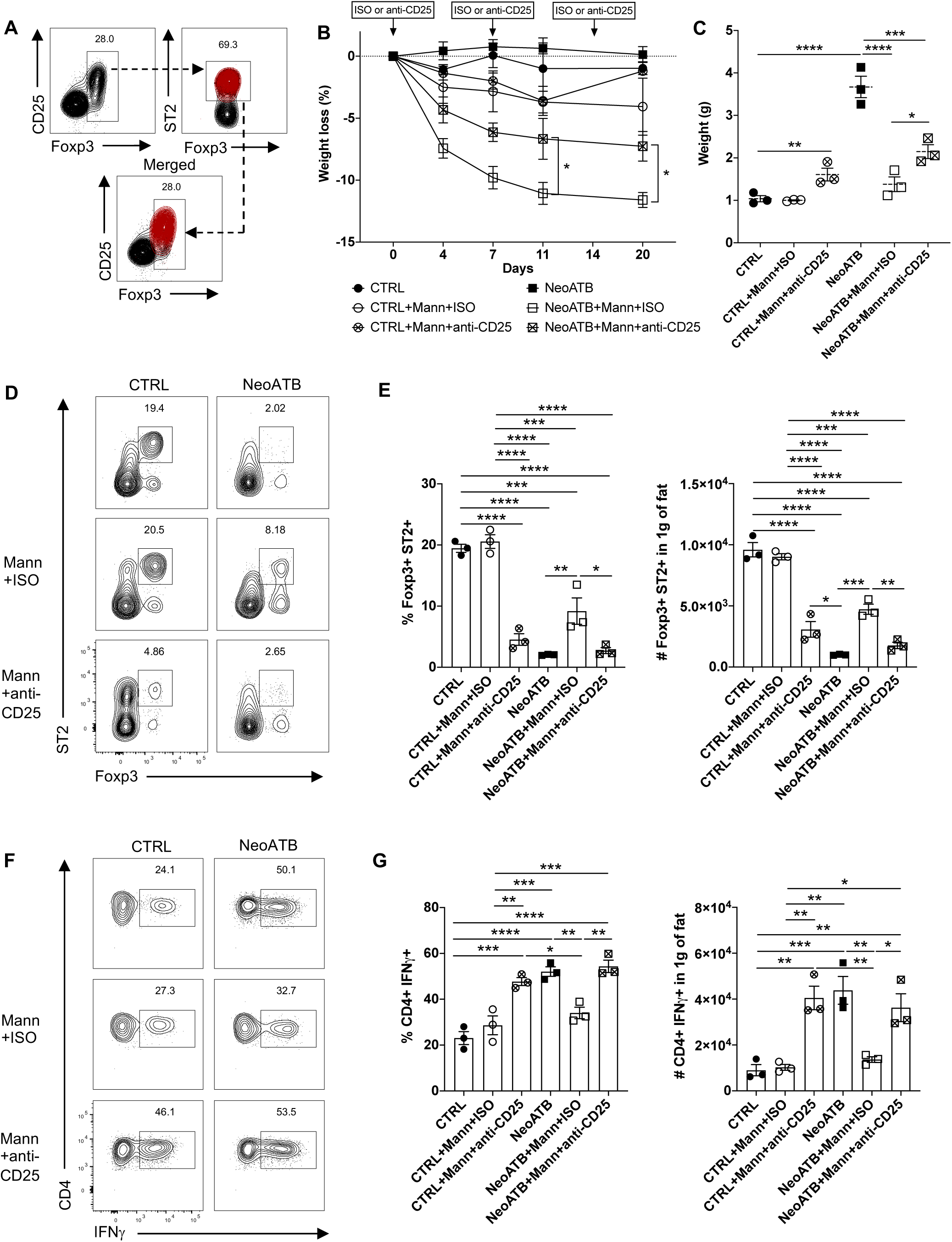
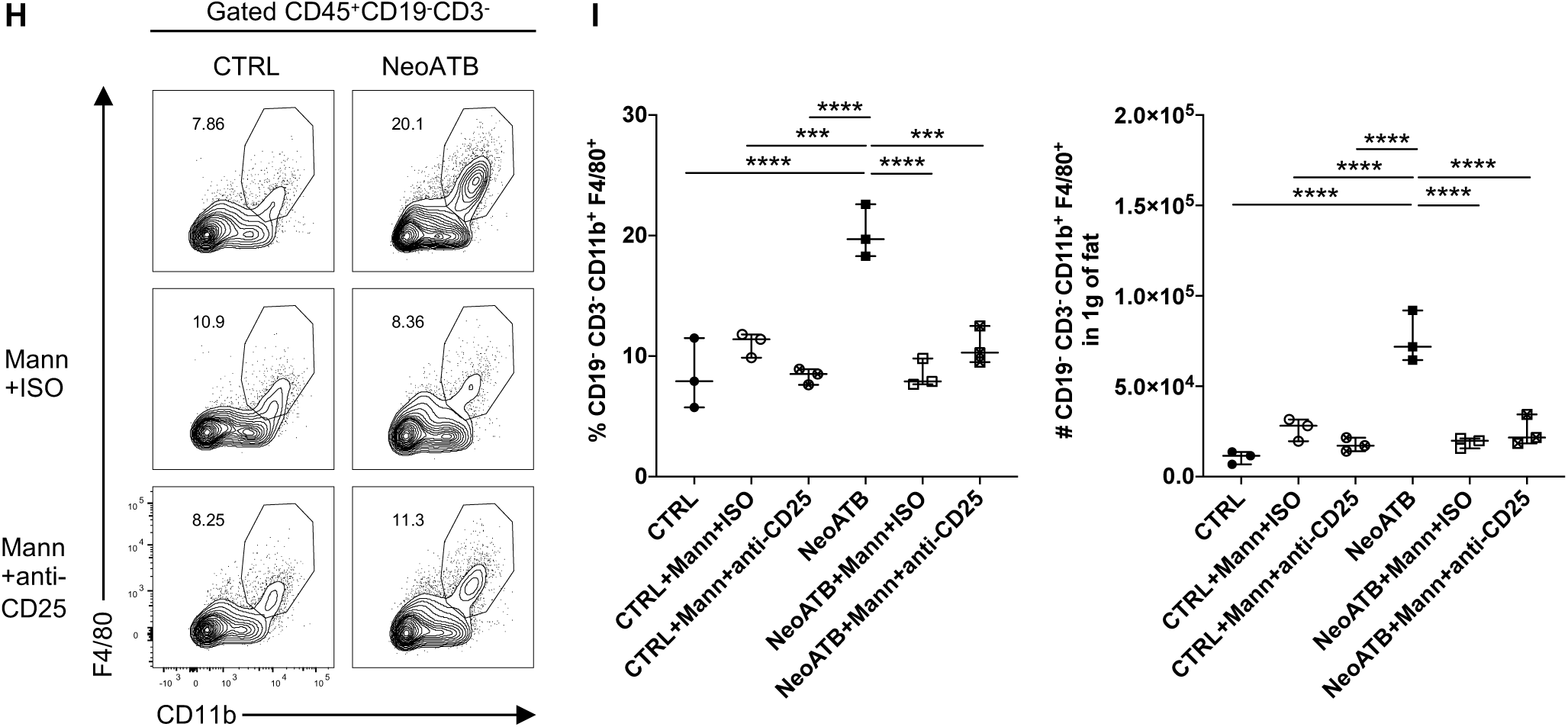
Treg depletion diminished the effect of D-mannose on the reduction of obesity in NeoATB mice. A. Representative FACS plot showing co-expression of CD25 and ST2 among adipose Foxp3^+^ cells. B. Adult aged (44-46 weeks old) CTRL and NeoATB mice were treated with/without D-Mannose or with Isotype or anti-CD25 neutralizing antibody. Weight loss (%) among different groups is shown. C. Weight of visceral adipose tissue measured at the end of experiment among different groups. D. Representative FACS plots showing frequency of adipose Treg cells (Gated as: CD45^+^TCRβ^+^CD4^+^Foxp3^+^ST2^+^) among different groups. E. Summarizing data showing frequency and total number of adipose Foxp3^+^ST2^+^ Treg cells among different groups. F. Representative FACS plots showing frequency of adipose Th1 IFNγ^+^ cells (Gated as: CD45^+^TCRβ^+^CD4^+^IFNγ^+^) among different groups. G. Summarizing data showing frequency and total number of adipose Th1 IFNγ^+^ cells among different groups. H. Representative FACS plots showing frequency of adipose macrophages (Gated as: CD45^+^CD19^-^CD3^-^CD11b^+^F4/80^+^) among different groups. I. Summarizing data showing frequency and total number of adipose infiltrating macrophages among different groups. Data are representative of 1 experiment (3 mice per cage/treatment). Statistical analysis was performed using one-way ANOVA (* p<0.05; ** p<0.01; *** p<0.001; **** p<0.0001). Statistical analysis in b) was performed using two-way ANOVA (* p<0.05). Gated: Live CD45^+^ Lin^-^

**Supplemental Figure 7.**
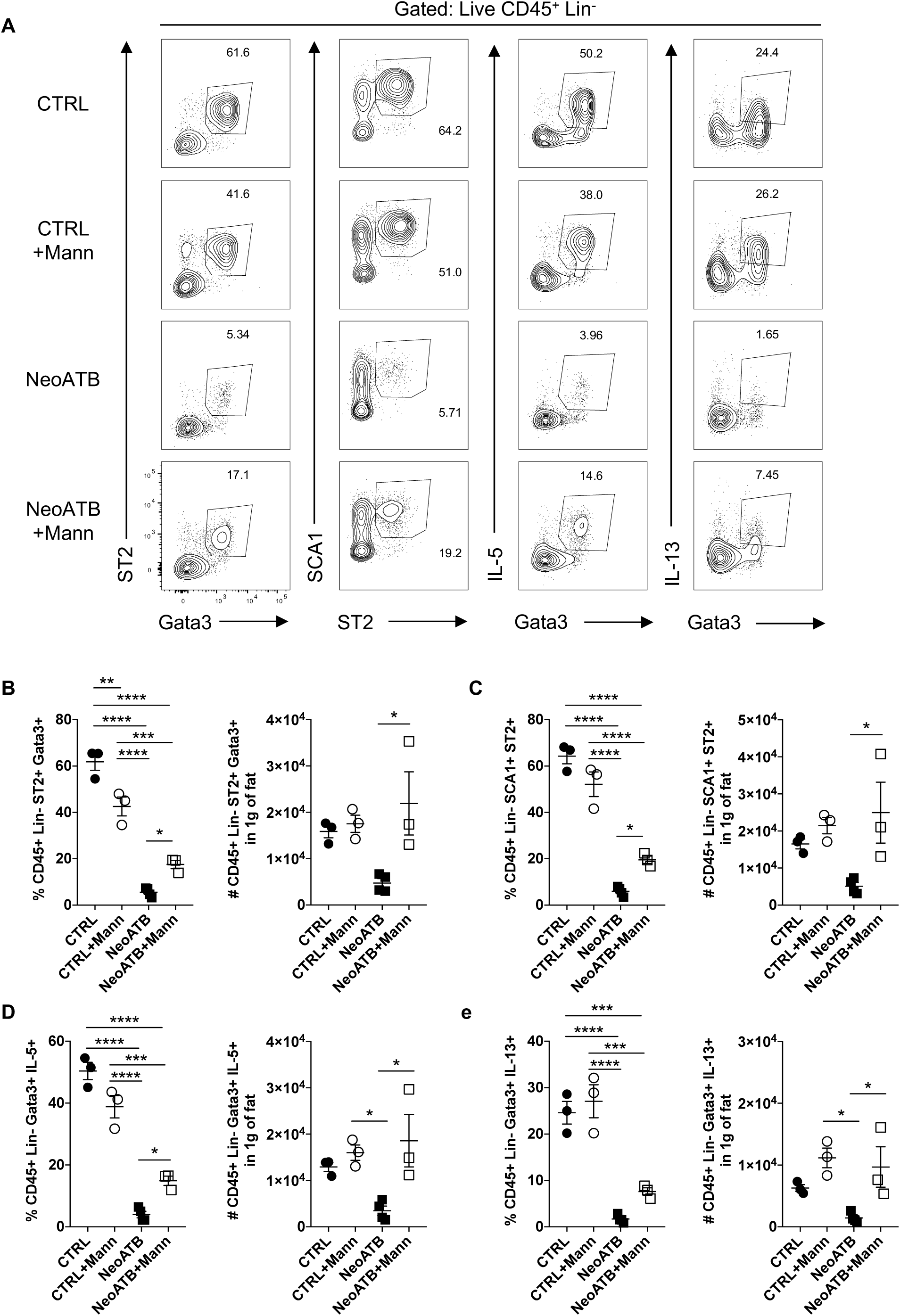
D-mannose increases adipose ILC2 cells in NeoATB mice. A. Representative FACS plots showing frequency of adipose ILC2 cells (Gated as: CD45^+^Lineage^-^ST2^+^Gata3^+^ or CD45^+^Lineage^-^ST2^+^SCA1^+^) among different groups. Representative FACS plots showing frequency of total IL-5^+^ or IL-13^+^ adipose ILC2 cells respectively (Gated as: CD45^+^Lineage^-^Gata3^+^IL-5^+^ or CD45^+^Lineage^-^Gata3^+^IL-13^+^). B. Summarizing data showing frequency and total number of adipose CD45^+^Lineage^-^ST2^+^Gata3^+^ ILC2 cells among different groups. C. Summarizing data showing frequency and total number of adipose CD45^+^Lineage^-^ST2^+^SCA1^+^ ILC2 cells among different groups. D. Summarizing data showing frequency and total number of adipose CD45^+^Lineage^-^Gata3^+^IL-5^+^ ILC2 cells among different groups. E. Summarizing data showing frequency and total number of adipose CD45^+^Lineage^-^Gata3^+^IL-13^+^ ILC2 cells among different groups. Data are representative of 1 out of 2 independent experiments (3-4 mice per cage/treatment). Statistical analysis was performed using one-way ANOVA (* p<0.05; ** p<0.01; *** p<0.001; **** p<0.0001).

**Supplemental Figure 8.**
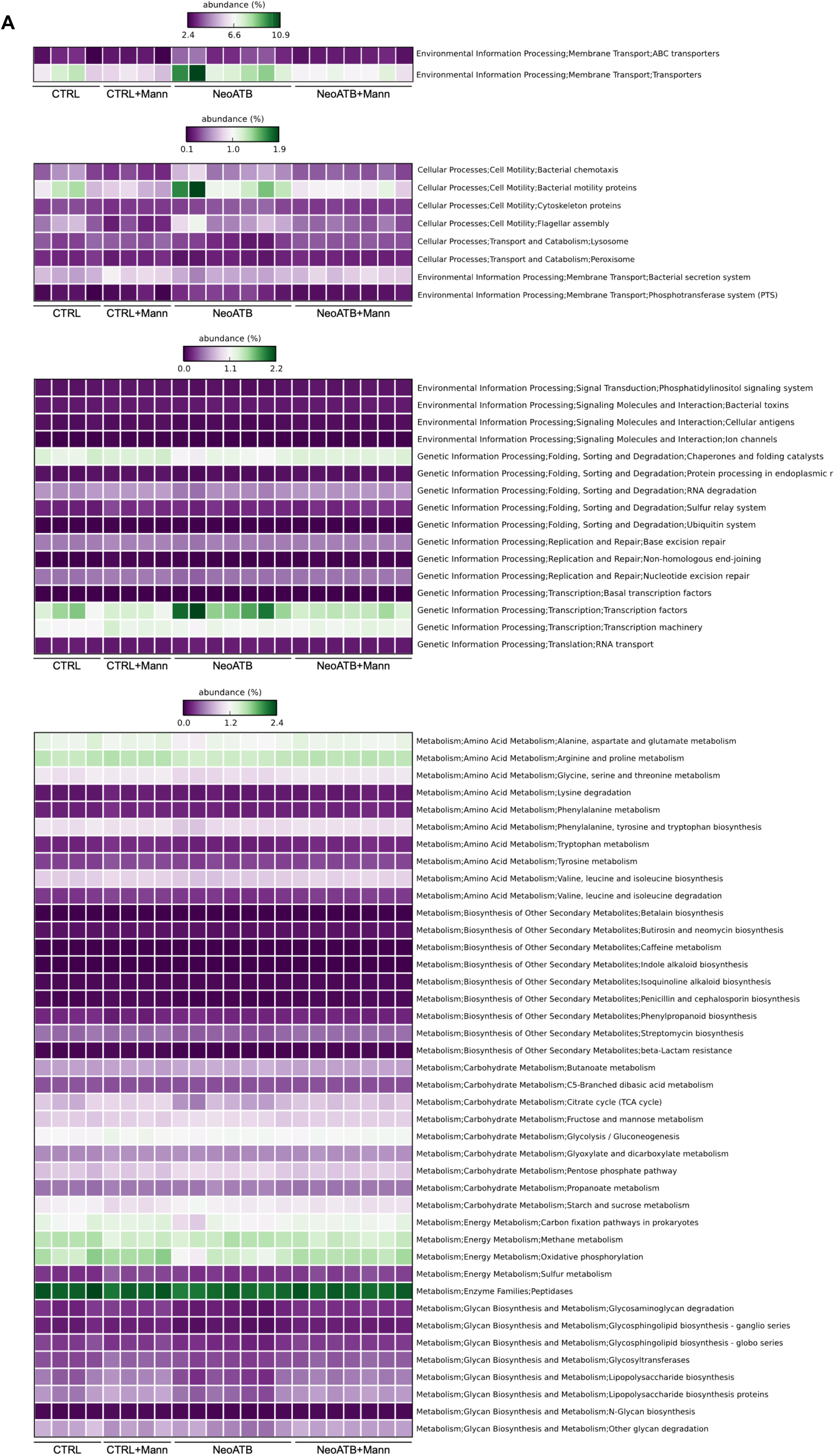
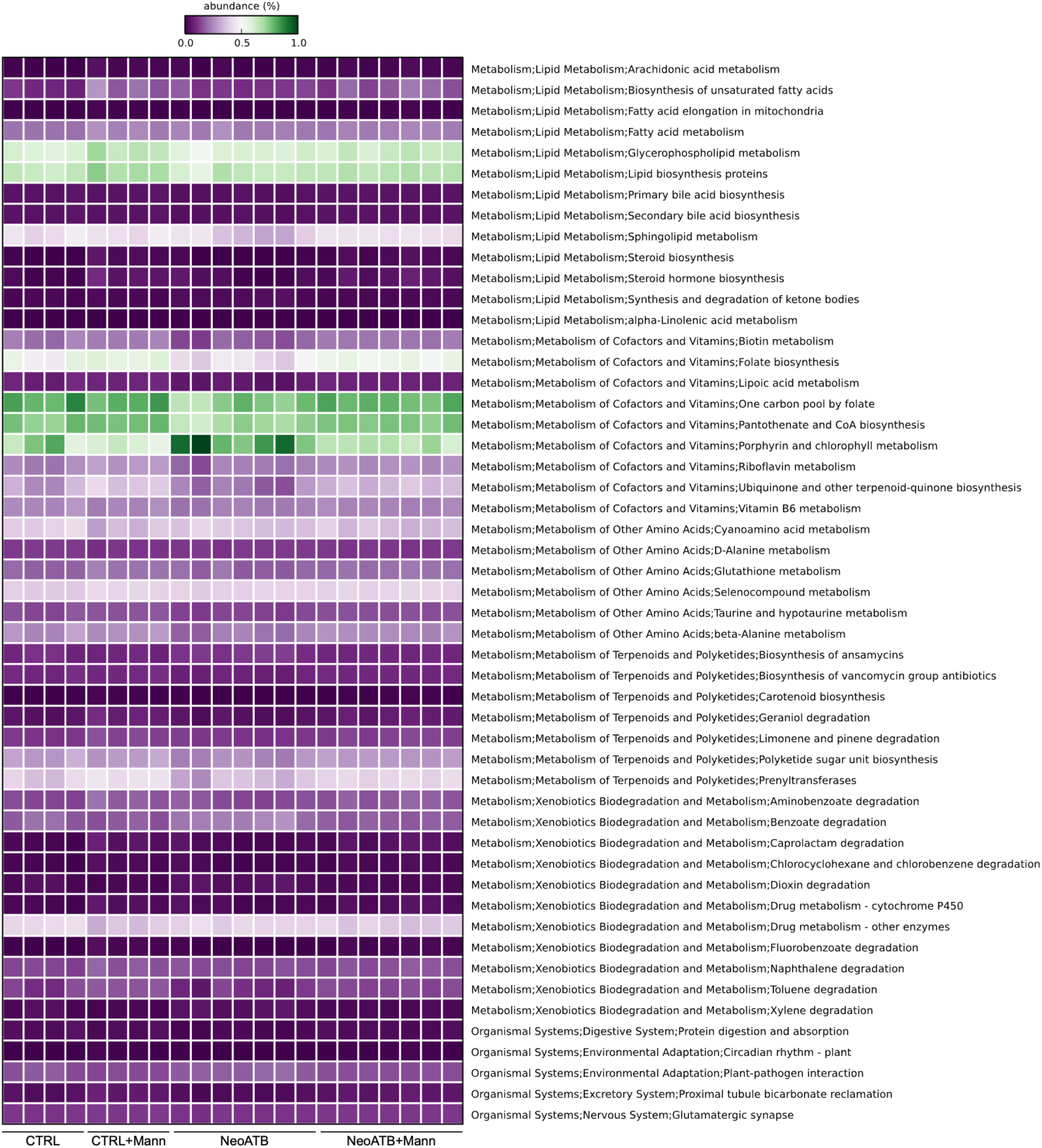
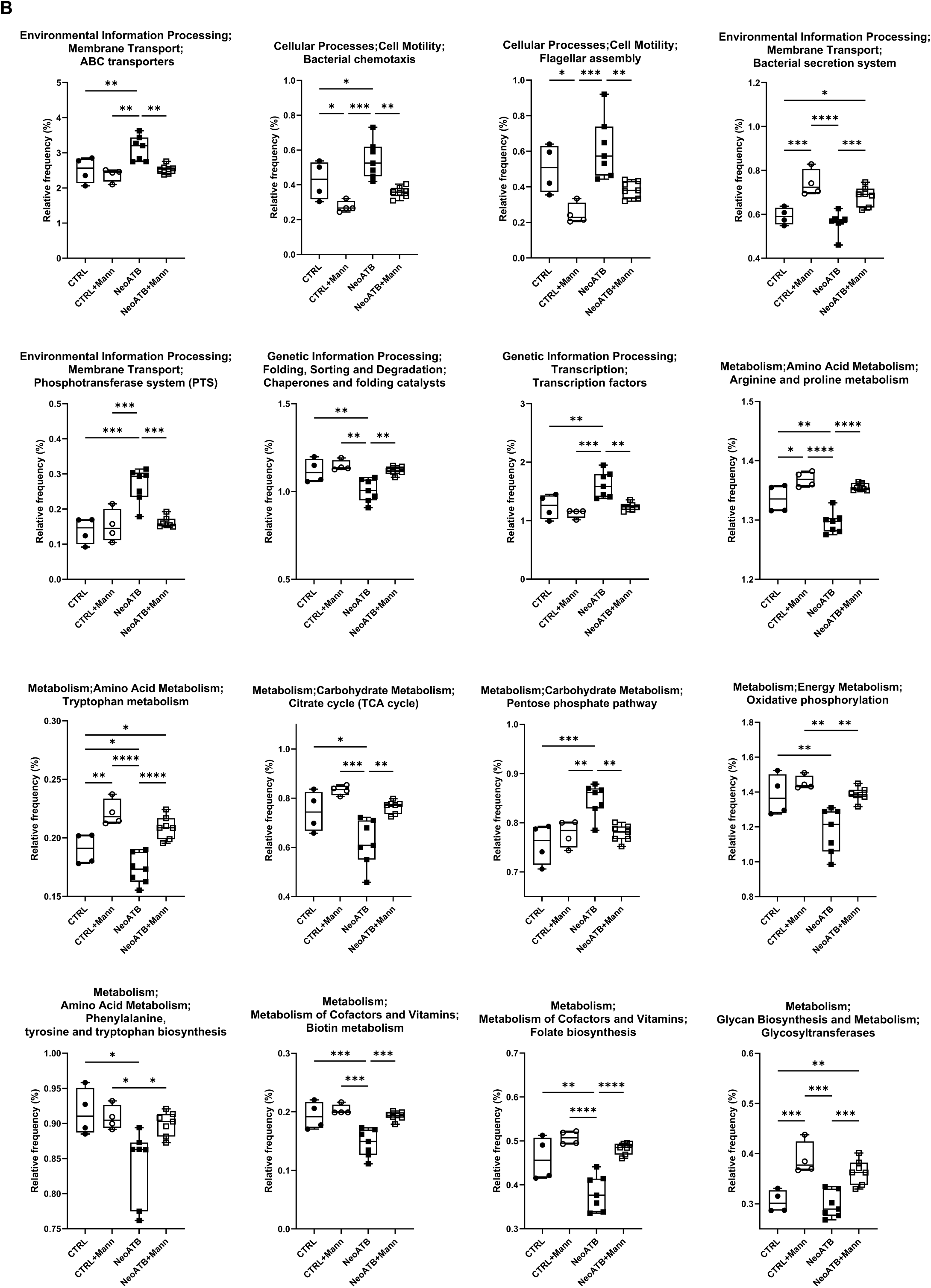
Comparative prediction of the functional metagenome of the gut microbiota. The figure shows a graphic representation of the significant predicted metabolic pathways using PICRUSt by analysis of the corresponding OTU table generated by QIIME for the bacterial communities. A. Heatmap plot showing all significant pathways (% of relative frequency of gene content prediction) among all groups identified by PICRUSt. B. Box-plot figures showing representative futures of the functional metagenome predictions. The Y-axis shows the relative frequencies of gene content prediction, and the X-axis shows the different groups. Significant differences were identified using One-way ANOVA test, followed by FDR (Benjamini Hochberg) posttest for multiple test correction.

**Supplemental Figure 9.**
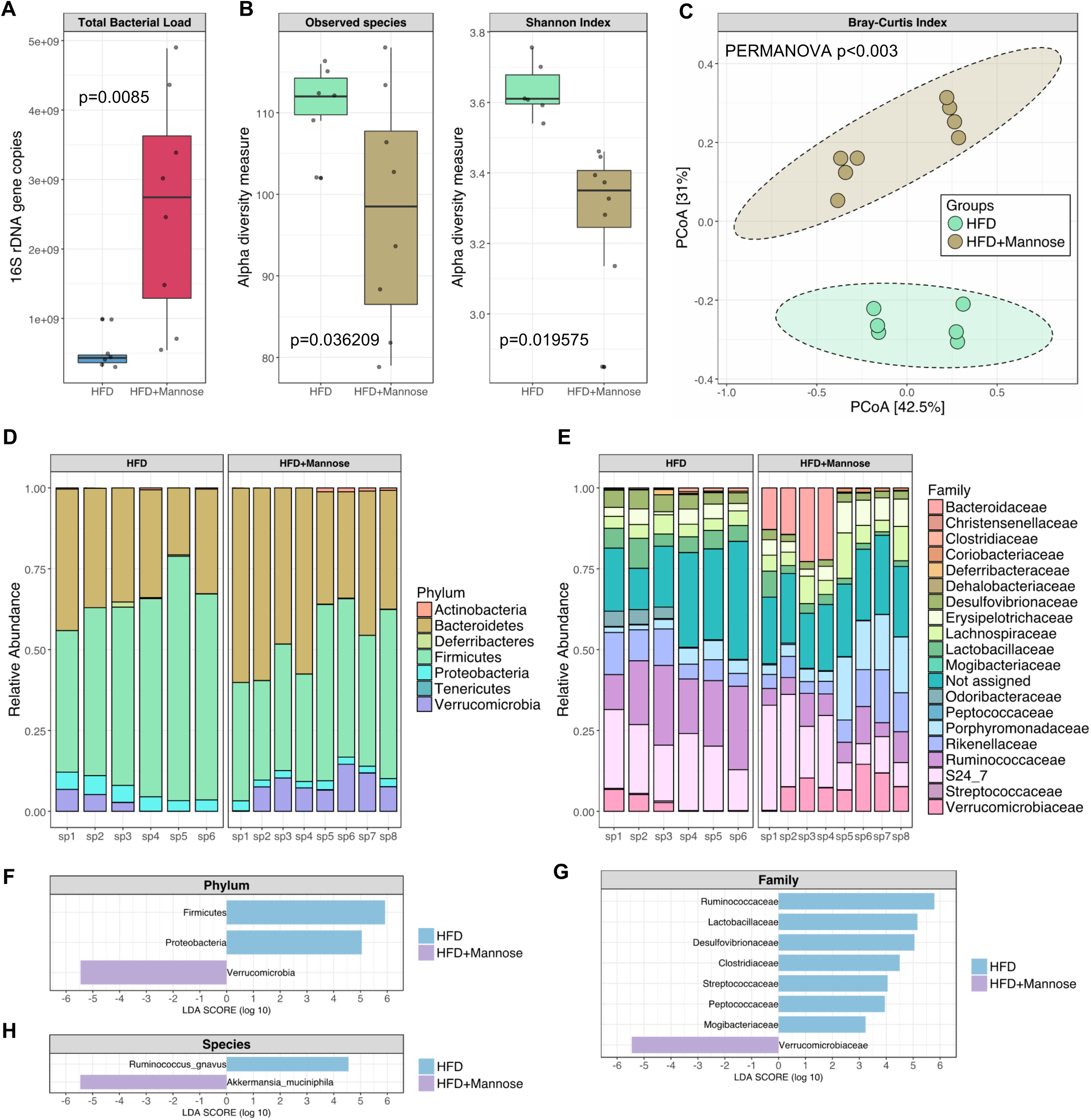
D-mannose corrects the microbiome of HFD treated mice. A. Total bacterial load determined as number of 16S rDNA gene copies using qPCR. B. Alpha diversity measure (Observed species, Shannon Index) between HFD and D-mannose treated HFD mice. C. Beta diversity measure (Bray-Curtis Index) between HFD and D-mannose treated HFD mice. D. Phylum level taxa abundance between HFD and D-mannose treated HFD mice. E. Family level taxa abundance between HFD and D-mannose treated HFD mice. F. Linear Discriminant Analysis (LDA=3) showing significantly different taxa on the phylum level determined by LEfSe analysis. G. Linear Discriminant Analysis (LDA=3) showing significantly different taxa on the family level determined by LEfSe analysis. H. Linear Discriminant Analysis (LDA=3) showing significantly different taxa on the species level determined by LEfSe analysis.

**Supplemental Figure 10.**
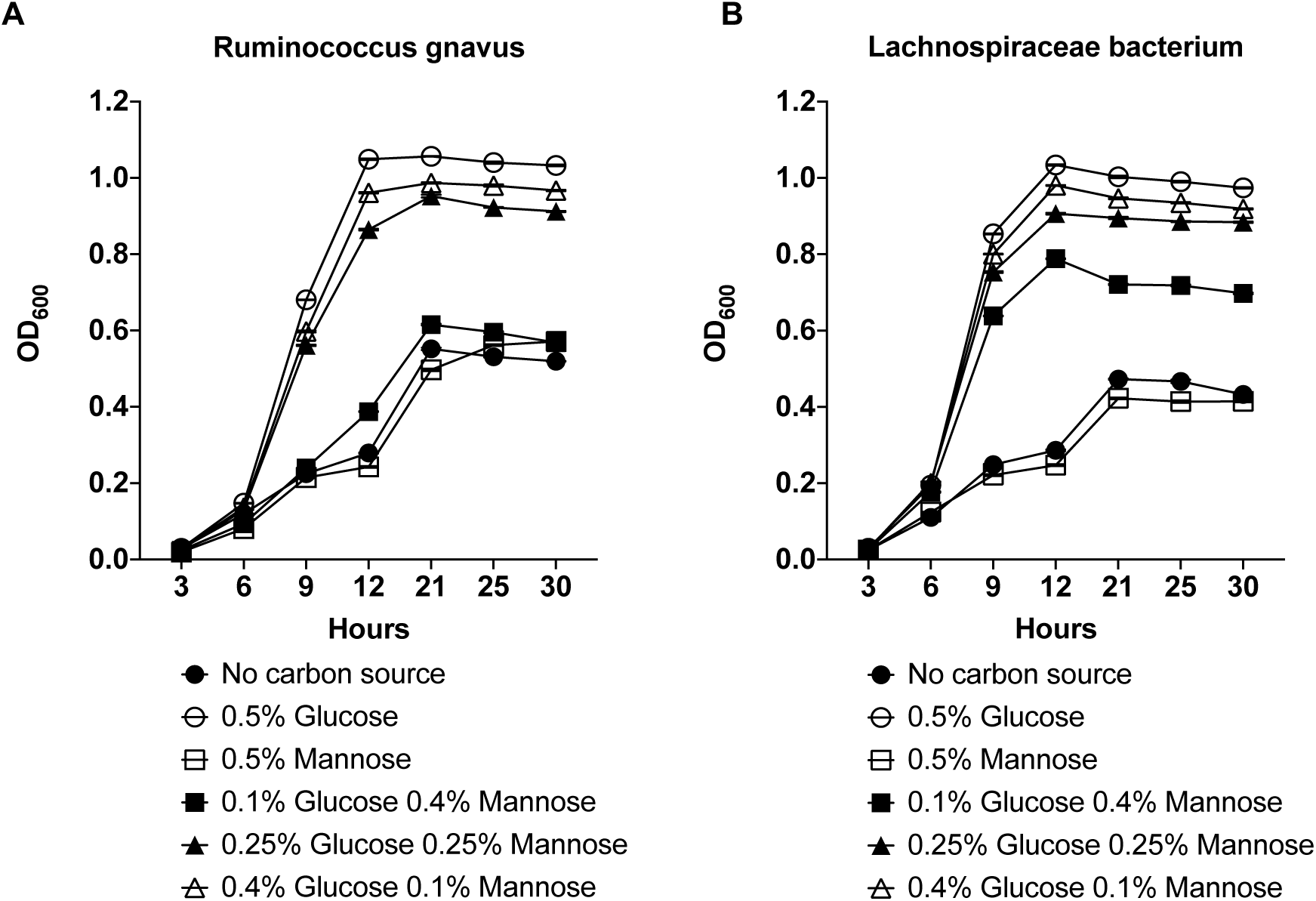
D-mannose suppresses growth of *Firmicutes*. A. Representative growth curve of *Ruminococcus gnavus* cultured in the heart infusion media without sugar (no carbon source) or different concentration ratios of D-Mannose : D-Glucose. B. Representative growth curve of *Lachnospiraceae bacterium* cultured in the heart infusion media without sugar (no carbon source) or different concentration ratios of D-Mannose : D-Glucose. Data are representative of one out of 3 independent experiments.

**Supplemental Table 1.** Antibody used for Flow cytometry.

| Antibody | Fluorochrome | Clone | Catalog number | Company |
| --- | --- | --- | --- | --- |
| Anti-mouse CD45 | Alexa Fluor 700 | 30-F11 | 56-0451-82 | Thermo Fisher Scientific |
| Anti-mouse CD4 | FITC | RM4-5 | 11-0042-85 | Thermo Fisher Scientific |
| Anti-mouse CD4 | Brilliant Violet 510 | RM4-5 | 100559 | Biolegend |
| Anti-mouse CD25 | PerCP-Cy5.5 | PC61.5 | 45-0251-82 | Thermo Fisher Scientific |
| Anti-mouse CD25 | eFluor660 | eBio7D4 | 50-0252-82 | Thermo Fisher Scientific |
| Anti-mouse TCRβ | APC-eFluor 780 | H57-597 | 47-5961-82 | Thermo Fisher Scientific |
| Anti-mouse Foxp3 | eFluor450 | FJK-16s | 48-5773-82 | Thermo Fisher Scientific |
| Anti-mouse ST2 | APC | RMST2-2 | 17-9335-82 | Thermo Fisher Scientific |
| Anti-mouse Ki67 | PE-eFluor610 | SolA15 | 61-5698-82 | Thermo Fisher Scientific |
| Anti-mouse TCR$\gamma\delta$ | PerCP-eFluor 710 | eBioGL3 | 46-5711-82 | Thermo Fisher Scientific |
| Anti-mouse IFNγ | eFluor450 | XMG1.2 | 48-7311-82 | Thermo Fisher Scientific |
| Anti-mouse IL-17A | APC | eBio17B7 | 17-7177-81 | Thermo Fisher Scientific |
| Anti-mouse CD8b | PE-Cy7 | eBioH35-17.2 | 25-0083-82 | Thermo Fisher Scientific |
| Anti-mouse Gata3 | PE-Cy7 | TWAJ | 25-9966-42 | Thermo Fisher Scientific |
| Anti-mouse SCA1 | FITC | D7 | 11-5981-81 | Thermo Fisher Scientific |
| Anti-mouse IL-5 | PE | TRFK5 | 12-7052-81 | Thermo Fisher Scientific |
| Anti-mouse IL-13 | PE-eFluor610 | eBio13A | 61-7133-82 | Thermo Fisher Scientific |
| Anti-mouse CD3e | APC-eFluor780 | 17A2 | 47-0032-82 | Thermo Fisher Scientific |
| Anti-mouse CD19 | PerCP-Cy5.5 | eBio1D3 | 45-0193-82 | Thermo Fisher Scientific |
| Anti-mouse CD11b | eFluor450 | M1/71 | 48-0112-82 | Thermo Fisher Scientific |
| Anti-mouse F4/80 | PE-eFluor610 | BM8 | 61-4801-82 | Invitrogen |
| Zombie Yellow™<br>Fixable Viability Kit | QDOT605 | N/A | 423104 | Biolegend |
| CD16/32Fc | N/A | 93 | 14-0161-86 | Thermo Fisher Scientific |

**Supplemental Table 2.**
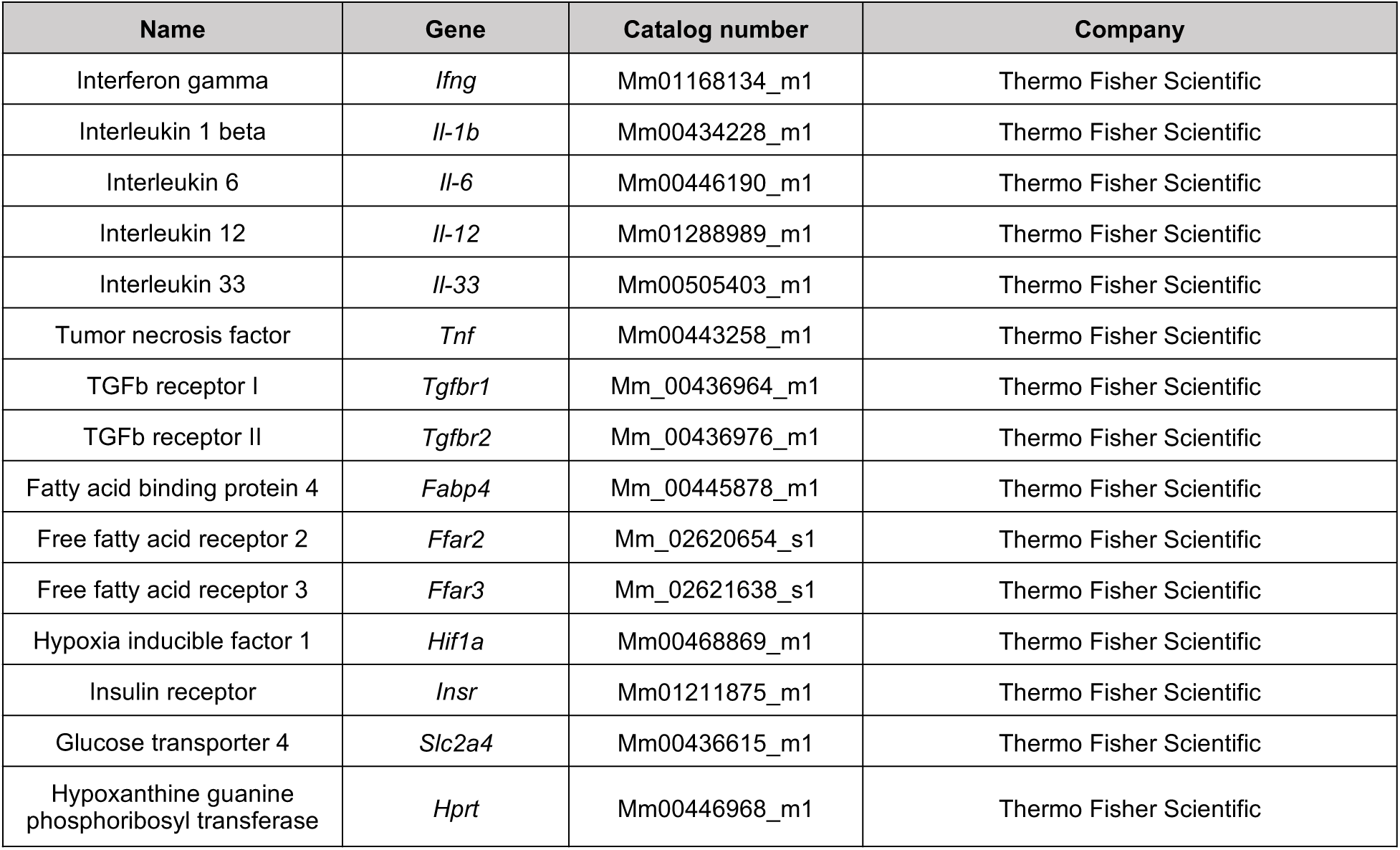
TaqMan Assays used for qPCR.

**Supplemental Table 3.** Antibody used for Western blot.

| Antibody | Catalog number | Dilution | Company |
| --- | --- | --- | --- |
| Insulin receptor alpha | bs-0047R-TR | 1:300 | Bioss Antibodies |
| Glut4 | bs-0384R-FR | 1:300 | Bioss Antibodies |
| HIF-1 Alpha | bs-0737R-TR | 1:200 | Bioss Antibodies |
| Anti-α-Tubulin | T5168-100UL | 1:50000 | Sigma |

**Supplemental Table 4.** Reagents used for Seahorse Assay.

| Reagent | Catalog number | Company |
| --- | --- | --- |
| Poly-L-Lysine | 25988-63-0 | MP biomedicals |
| RPMI medium | 103576-100 | Agilent technologies |
| Glucose | G6152-1kg | Sigma |
| L-Glutamine | 25030164 | Gibco |
| Pyruvate | 13-115E | Lonza |
| Oligomycin | 75351-5mg | Sigma |
| CCCP | C2920-10mg | Sigma |
| Rotenone | R8875-1g | Sigma |
| Antimycin A | A8674-25mg | Sigma |

**Supplemental Table 5.** Instruments.

| Instrument | Company |
| --- | --- |
| BD LSRFortessa | BD Biosciences |
| BD FACSAria III Cell Sorter | BD Biosciences |
| Leica Aperio Scan Scope | Leica Biosystems |
| Seahorse XFe96 Analyzer | Agilent |
| SpectraMax Plus 384 Microplate Reader | Molecular devices |
| Accu-Chek Aviva Blood Glucometer | Accu Chek |
| Whitley A35 Workstation | Don Whitley Scientific |
| Applied Biosystems 7500 Real Time PCR System | Applied Biosystems by Thermo Fisher Scientific |
| QuantStudio 3 | Applied Biosystems by Thermo Fisher Scientific |

**Supplemental Table 6.** Software.

| Software | Company | Web |
| --- | --- | --- |
| FlowJo 10 | BD Biosciences | https://www.flowjo.com/solutions/flowjo |
| Prism 8 | GraphPad | https://www.graphpad.com/scientific-software/prism/ |
| Aperio Image Scope | Leica Biosystems | https://www.leicabiosystems.com/digital-pathology/manage/aperio-imagescope/ |
| AdipoCount | n/a | http://www.csbio.sjtu.edu.cn/bioinf/AdipoCount/ |
| STAMP v2.1.3 | n/a | https://beikolab.cs.dal.ca/software/STAMP |

## References

1. Bluher M. Obesity: global epidemiology and pathogenesis. Nat Rev Endocrinol. 2019;15(5):288–98.

2. Khan MT, Nieuwdorp M, and Backhed F. Microbial modulation of insulin sensitivity. Cell Metab. 2014;20(5):753–60.

3. Ridaura VK, Faith JJ, Rey FE, Cheng J, Duncan AE, Kau AL, et al. Gut microbiota from twins discordant for obesity modulate metabolism in mice. Science. 2013;341(6150):1241214.

4. Turnbaugh PJ, Ley RE, Mahowald MA, Magrini V, Mardis ER, and Gordon JI. An obesity-associated gut microbiome with increased capacity for energy harvest. Nature. 2006;444(7122):1027–31.

5. Ajslev TA, Andersen CS, Gamborg M, Sorensen TI, and Jess T. Childhood overweight after establishment of the gut microbiota: the role of delivery mode, pre-pregnancy weight and early administration of antibiotics. Int J Obes (Lond). 2011;35(4):522–9.

6. Murphy R, Stewart AW, Braithwaite I, Beasley R, Hancox RJ, Mitchell EA, et al. Antibiotic treatment during infancy and increased body mass index in boys: an international cross-sectional study. Int J Obes (Lond). 2014;38(8):1115–9.

7. Dominguez-Bello MG, Costello EK, Contreras M, Magris M, Hidalgo G, Fierer N, et al. Delivery mode shapes the acquisition and structure of the initial microbiota across multiple body habitats in newborns. Proc Natl Acad Sci U S A. 2010;107(26):11971–5.

8. Blustein J, Attina T, Liu M, Ryan AM, Cox LM, Blaser MJ, et al. Association of caesarean delivery with child adiposity from age 6 weeks to 15 years. Int J Obes (Lond). 2013;37(7):900–6.

9. Zhang D, Chia C, Jiao X, Jin W, Kasagi S, Wu R, et al. D-mannose induces regulatory T cells and suppresses immunopathology. Nat Med. 2017;23(9):1036–45.

10. Fabbrini E, Sullivan S, and Klein S. Obesity and nonalcoholic fatty liver disease: biochemical, metabolic, and clinical implications. Hepatology. 2010;51(2):679–89.

11. Petersen MC, and Shulman GI. Mechanisms of Insulin Action and Insulin Resistance. Physiol Rev. 2018;98(4):2133–223.

12. Leto D, and Saltiel AR. Regulation of glucose transport by insulin: traffic control of GLUT4. Nat Rev Mol Cell Biol. 2012;13(6):383–96.

13. Jager J, Gremeaux T, Cormont M, Le Marchand-Brustel Y, and Tanti JF. Interleukin-1beta-induced insulin resistance in adipocytes through down-regulation of insulin receptor substrate-1 expression. Endocrinology. 2007;148(1):241–51.

14. Stephens JM, Lee J, and Pilch PF. Tumor necrosis factor-alpha-induced insulin resistance in 3T3-L1 adipocytes is accompanied by a loss of insulin receptor substrate-1 and GLUT4 expression without a loss of insulin receptor-mediated signal transduction. J Biol Chem. 1997;272(2):971–6.

15. Wada T, Hoshino M, Kimura Y, Ojima M, Nakano T, Koya D, et al. Both type I and II IFN induce insulin resistance by inducing different isoforms of SOCS expression in 3T3-L1 adipocytes. Am J Physiol Endocrinol Metab. 2011;300(6):E1112–23.

16. Wentworth JM, Zhang JG, Bandala-Sanchez E, Naselli G, Liu R, Ritchie M, et al. Interferon-gamma released from omental adipose tissue of insulin-resistant humans alters adipocyte phenotype and impairs response to insulin and adiponectin release. Int J Obes (Lond). 2017;41(12):1782–9.

17. Hosogai N, Fukuhara A, Oshima K, Miyata Y, Tanaka S, Segawa K, et al. Adipose tissue hypoxia in obesity and its impact on adipocytokine dysregulation. Diabetes. 2007;56(4):901–11.

18. Ye J, Gao Z, Yin J, and He Q. Hypoxia is a potential risk factor for chronic inflammation and adiponectin reduction in adipose tissue of ob/ob and dietary obese mice. Am J Physiol Endocrinol Metab. 2007;293(4):E1118–28.

19. Feuerer M, Herrero L, Cipolletta D, Naaz A, Wong J, Nayer A, et al. Lean, but not obese, fat is enriched for a unique population of regulatory T cells that affect metabolic parameters. Nat Med. 2009;15(8):930–9.

20. Vasanthakumar A, Moro K, Xin A, Liao Y, Gloury R, Kawamoto S, et al. The transcriptional regulators IRF4, BATF and IL-33 orchestrate development and maintenance of adipose tissue-resident regulatory T cells. Nat Immunol. 2015;16(3):276–85.

21. Tu E, Chia CPZ, Chen W, Zhang D, Park SA, Jin W, et al. T Cell Receptor-Regulated TGF-beta Type I Receptor Expression Determines T Cell Quiescence and Activation. Immunity. 2018;48(4):745–59 e6.

22. Brestoff JR, Kim BS, Saenz SA, Stine RR, Monticelli LA, Sonnenberg GF, et al. Group 2 innate lymphoid cells promote beiging of white adipose tissue and limit obesity. Nature. 2015;519(7542):242–6.

23. Walker JA, and McKenzie AN. Development and function of group 2 innate lymphoid cells. Curr Opin Immunol. 2013;25(2):148–55.

24. Backhed F, Ding H, Wang T, Hooper LV, Koh GY, Nagy A, et al. The gut microbiota as an environmental factor that regulates fat storage. P Natl Acad Sci USA. 2004;101(44):15718–23.

25. Ley RE, Turnbaugh PJ, Klein S, and Gordon JI. Microbial ecology: human gut microbes associated with obesity. Nature. 2006;444(7122):1022–3.

26. Turnbaugh PJ, Backhed F, Fulton L, and Gordon JI. Diet-induced obesity is linked to marked but reversible alterations in the mouse distal gut microbiome. Cell Host Microbe. 2008;3(4):213–23.

27. Langille MG, Zaneveld J, Caporaso JG, McDonald D, Knights D, Reyes JA, et al. Predictive functional profiling of microbial communities using 16S rRNA marker gene sequences. Nat Biotechnol. 2013;31(9):814–21.

28. Parks DH, Tyson GW, Hugenholtz P, and Beiko RG. STAMP: statistical analysis of taxonomic and functional profiles. Bioinformatics. 2014;30(21):3123–4.

29. Murphy EF, Cotter PD, Healy S, Marques TM, O’Sullivan O, Fouhy F, et al. Composition and energy harvesting capacity of the gut microbiota: relationship to diet, obesity and time in mouse models. Gut. 2010;59(12):1635–42.

30. Fernandes J, Su W, Rahat-Rozenbloom S, Wolever TM, and Comelli EM. Adiposity, gut microbiota and faecal short chain fatty acids are linked in adult humans. Nutr Diabetes. 2014;4:e121.

31. Schwiertz A, Taras D, Schafer K, Beijer S, Bos NA, Donus C, et al. Microbiota and SCFA in lean and overweight healthy subjects. Obesity (Silver Spring). 2010;18(1):190–5.

32. Tirosh A, Calay ES, Tuncman G, Claiborn KC, Inouye KE, Eguchi K, et al. The short-chain fatty acid propionate increases glucagon and FABP4 production, impairing insulin action in mice and humans. Sci Transl Med. 2019;11(489).

33. Canfora EE, Jocken JW, and Blaak EE. Short-chain fatty acids in control of body weight and insulin sensitivity. Nat Rev Endocrinol. 2015;11(10):577–91.

34. Goossens GH, and Blaak EE. Adipose tissue dysfunction and impaired metabolic health in human obesity: a matter of oxygen? Front Endocrinol (Lausanne). 2015;6:55.

35. Rausch ME, Weisberg S, Vardhana P, and Tortoriello DV. Obesity in C57BL/6J mice is characterized by adipose tissue hypoxia and cytotoxic T-cell infiltration. Int J Obes (Lond). 2008;32(3):451–63.

36. Miller AM, Asquith DL, Hueber AJ, Anderson LA, Holmes WM, McKenzie AN, et al. Interleukin-33 induces protective effects in adipose tissue inflammation during obesity in mice. Circ Res. 2010;107(5):650–8.

37. Torretta S, Scagliola A, Ricci L, Mainini F, Di Marco S, Cuccovillo I, et al. D-mannose suppresses macrophage IL-1beta production. Nat Commun. 2020;11(1):6343.

38. Ley RE, Backhed F, Turnbaugh P, Lozupone CA, Knight RD, and Gordon JI. Obesity alters gut microbial ecology. Proc Natl Acad Sci U S A. 2005;102(31):11070–5.

39. Sharma V, Smolin J, Nayak J, Ayala JE, Scott DA, Peterson SN, et al. Mannose Alters Gut Microbiome, Prevents Diet-Induced Obesity, and Improves Host Metabolism. Cell Rep. 2018;24(12):3087–98.

40. Riva A, Borgo F, Lassandro C, Verduci E, Morace G, Borghi E, et al. Pediatric obesity is associated with an altered gut microbiota and discordant shifts in Firmicutes populations. Environ Microbiol. 2017;19(1):95–105.

41. Hou YP, He QQ, Ouyang HM, Peng HS, Wang Q, Li J, et al. Human Gut Microbiota Associated with Obesity in Chinese Children and Adolescents. Biomed Res Int. 2017;2017:7585989.

42. Louis S, Tappu RM, Damms-Machado A, Huson DH, and Bischoff SC. Characterization of the Gut Microbial Community of Obese Patients Following a Weight-Loss Intervention Using Whole Metagenome Shotgun Sequencing. PLoS One. 2016;11(2):e0149564.

43. Cussotto S, Delgado I, Anesi A, Dexpert S, Aubert A, Beau C, et al. Tryptophan Metabolic Pathways Are Altered in Obesity and Are Associated With Systemic Inflammation. Front Immunol. 2020;11:557.

44. Del Chierico F, Abbatini F, Russo A, Quagliariello A, Reddel S, Capoccia D, et al. Gut Microbiota Markers in Obese Adolescent and Adult Patients: Age-Dependent Differential Patterns. Front Microbiol. 2018;9:1210.

45. Kose S, Sozlu S, Bolukbasi H, Unsal N, and Gezmen-Karadag M. Obesity is associated with folate metabolism. Int J Vitam Nutr Res. 2020;90(3-4):353–64.

46. Via M. The malnutrition of obesity: micronutrient deficiencies that promote diabetes. ISRN Endocrinol. 2012;2012:103472.

47. Cani PD, and de Vos WM. Next-Generation Beneficial Microbes: The Case of Akkermansia muciniphila. Front Microbiol. 2017;8:1765.

48. Depommier C, Everard A, Druart C, Plovier H, Van Hul M, Vieira-Silva S, et al. Supplementation with Akkermansia muciniphila in overweight and obese human volunteers: a proof-of-concept exploratory study. Nat Med. 2019;25(7):1096–103.

49. Cirera S. Highly efficient method for isolation of total RNA from adipose tissue. BMC Res Notes. 2013;6:472.

50. Prochazkova M, Hall B, Hu M, Okine T, Reukauf J, Binukumar BK, et al. Peripheral and orofacial pain sensation is unaffected by the loss of p39. Mol Pain. 2017;13:1744806917737205.

51. Dunham-Snary KJ, Sandel MW, Westbrook DG, and Ballinger SW. A method for assessing mitochondrial bioenergetics in whole white adipose tissues. Redox Biol. 2014;2:656–60.

52. Weber N, Liou D, Dommer J, MacMenamin P, Quinones M, Misner I, et al. Nephele: a cloud platform for simplified, standardized and reproducible microbiome data analysis. Bioinformatics. 2018;34(8):1411–3.

53. Dhariwal A, Chong J, Habib S, King IL, Agellon LB, and Xia J. MicrobiomeAnalyst: a web-based tool for comprehensive statistical, visual and meta-analysis of microbiome data. Nucleic Acids Res. 2017;45(W1):W180–W8.

54. R-Core-Team. R: A language and environment for statistical computing. R Foundation for Statistical Computing, Vienna, Austria. 2013.

55. Zhi X, Wang J, Lu P, Jia J, Shen HB, and Ning G. AdipoCount: A New Software for Automatic Adipocyte Counting. Front Physiol. 2018;9:85.

56. Liang W, Menke AL, Driessen A, Koek GH, Lindeman JH, Stoop R, et al. Establishment of a general NAFLD scoring system for rodent models and comparison to human liver pathology. PLoS One. 2014;9(12):e115922.

